# ZNF687 couples bone marrow myeloid progenitor dynamics to osteoclastogenesis in severe Paget’s disease of bone

**DOI:** 10.64898/2026.08.11.744220

**Authors:** Sharon Russo, Vincenzo Lullo, Alessandro Miranda, Dario Acampora, Danilo Licastro, Maria Strazzullo, Carmine Settembre, Maria Rosaria Matarazzo, Antonio Simeone, Fernando Gianfrancesco

## Abstract

Paget’s disease of bone (PDB) is a late-onset skeletal disorder characterized by excessive osteoclast-mediated bone remodelling and disorganized bone deposition. The P937R mutation in the *ZNF687* gene causes a severe form of PDB complicated by giant cell tumour transformation. Although ZNF687 has been implicated in osteoclastogenesis, whether it regulates upstream haematopoietic progenitor dynamics and bone marrow myeloid output remains unclear.

Using a constitutive *Zfp687* knock-out mouse model, we showed that *Zfp687* loss causes postnatal growth restriction, reduced bone marrow cellularity, impaired osteoclast differentiation *in vitro* and *in vivo*, and increased trabecular bone mass during adulthood. Flow cytometry revealed a marked reduction in osteoclast progenitors and macrophages in *Zfp687*-deficient bone marrow, whereas the pagetic P937R mutation promoted the expansion of the same myeloid populations in the *Zfp687*^P937R^ knock-in mouse model. Single-cell RNA sequencing of bone marrow-derived c-Kit^+^ haematopoietic progenitors further demonstrated that *Zfp687* loss selectively disrupted the myeloid progenitor compartment. This analysis identified 22 transcriptionally distinct populations and revealed a significant depletion of the early cycling granulocyte–monocyte progenitor cluster, without evidence of a global block in myeloid differentiation. Mechanistically, *Zfp687* deficiency impaired the Brd4-c-Myc-NFATc1 axis in osteoclastogenic precursors and reduced *Csf1* expression in bone marrow stromal and osteoblastic cells, linking intrinsic transcriptional competence to niche-derived M-CSF support. In pagetic patient iPSCs-derived haematopoietic progenitors, the P937R mutation enhanced clonogenic haematopoietic output, accelerated colony formation, and promoted the expansion of primitive/multipotent colony-forming progenitors, leading to hypercellular myeloid colonies.

Together, our findings establish ZNF687 as a regulator of haematopoietic progenitor dynamics that couples bone marrow myeloid output to osteoclastogenesis, providing a progenitor-level mechanism for severe ZNF687-related PDB.

## Introduction

Paget’s Disease of Bone (PDB) is a chronic skeletal disorder characterized by excessive and disorganized bone remodelling, resulting in bone pain, deformities, and fractures^1,2^. PDB has traditionally been described as a disorder of abnormal osteoclast activity, in which hyperactive and multinucleated osteoclasts drive excessive bone resorption followed by disorganized compensatory bone formation^3^. However, osteoclasts originate from the bone marrow (BM) resident myeloid progenitors and monocyte/macrophage precursors suggesting that alterations occurring upstream of terminal osteoclast differentiation may also contribute to pathologic bone remodelling^4^. Despite the recognized role of osteoclast precursors in skeletal diseases, the mechanisms linking PDB-associated genetic defects to the regulation of haematopoietic progenitor dynamics remain largely unexplored.

Osteoclast differentiation depends on coordinated expansion, survival, and commitment of myeloid progenitors within the BM microenvironment. Among these populations, granulocyte-monocyte progenitors (GMPs), common monocyte progenitors (cMoP), and downstream macrophage-lineage cells represent critical sources of osteoclast-competent precursors^5–7^. Their differentiation into mature, multi-nucleated osteoclasts requires the macrophages colony stimulating factor (M-CSF) and the receptor activator of nuclear factor κB (NFκB) ligand (RANKL), which together support precursors proliferation, survival, fusion, and activation^8–13^. While the molecular mechanisms controlling terminal osteoclast differentiation have been extensively investigated, significantly less is known about how the abundance, proliferative state, and lineage competence of the osteoclast progenitor pool are established and maintained in physiological and pathological contexts^14–16^.

ZNF687 is a transcription factor linked to a severe and rare form of PDB^17–19^. The P937R mutation in *ZNF687* is associated with an aggressive PDB phenotype, characterized by early disease onset, polyostotic pattern, increased risk of giant cell tumour transformation, and reduced responsiveness to the standard anti-resorptive therapies^18,19^. Previous studies have demonstrated that ZNF687 expression increases during peripheral blood mononuclear cells differentiation into osteoclasts, with higher expression levels detected in presence of the P937R mutation^20^. Moreover, the *Zfp687*^P937R^ knock-in (KI) mouse model displayed enhanced osteoclastogenesis *in vivo*, whereas engineered *Zfp687*^+/−^ knock-out (KO) RAW264.7 cells showed impaired osteoclast differentiation upon RANKL stimulation^20^. These findings clearly highlight a key role for ZNF687 in osteoclast biology, but whether ZNF687 also regulates haematopoietic progenitors states that supply the osteoclastogenic pool remains unknown.

A potential molecular mediator of this dual function is bromodomain-containing protein 4 (BRD4), a BET-family chromatin reader involved in transcriptional control, lineage specification, and cellular differentiation^21,22^. BRD4 plays important roles in stem and progenitor cell fate decisions, and osteoclast differentiation^23–25^. In this context, BRD4 regulates macrophage development and supports RANKL-induced transcriptional programmes, including c-Myc and NFATc1 activation^26–28^. Beyond bone, BRD4 is also deeply involved in hematopoietic lineage specification and commitment^29–32^. Notably, a direct interaction between znf687 and brd4 has been described in Zebrafish during late-stage neutrophil differentiation, suggesting a potential functional connection between ZNF687, BRD4-dependent transcription, and myeloid lineage control^32^. However, whether ZNF687 regulates BRD4-associated programmes in BM-derived myeloid progenitor and osteoclastogenic precursors remains unclear.

In this study, we investigated the role of ZNF687 in BM haematopoietic and osteoclastogenic remodelling, using a constitutive *Zfp687*-KO and the *Zfp687*^P937R^-KI mouse models, single-cell transcriptomic profiling of BM-derived haematopoietic progenitors, and human iPSC-derived haematopoietic differentiation using cells from a P937R-mutant patient and a matched healthy sibling control. We show that ZNF687 regulates BM myeloid progenitor dynamics, with *Zfp687* loss causing depletion of the cycling GMP population, reduction of osteoclast progenitors and macrophages, impaired osteoclastogenesis, and increased trabecular bone mass. Conversely, the P937R mutation expands BM myeloid populations *in vivo* and enhances clonogenic haematopoietic output in human iPSC-derived progenitors.

Together, our findings identify ZNF687 as a regulator of bone marrow myeloid progenitor dynamics and stromal niche support, providing a progenitor-level mechanism for the excessive osteoclastogenic remodelling characteristic of severe ZNF687-related PDB.

## Results

### Zfp687 loss impairs postnatal skeletal growth and marrow cavity development

The postnatal maturation of the BM compartment requires coordinated skeletal remodelling, vascular invasion, and osteoclast-dependent expansion of the medullary cavity, thereby linking osteoclast activity to the establishment and maintenance of the haematopoietic microenvironment^33^. To investigate the role of ZNF687 in myeloid BM-derived cells during bone remodelling, and to assess the functional consequences of its loss relative to the activating P937R mutation, we generated a constitutive *Zfp687* knock-out mouse model (here referred to as *Zfp687*^−/−^) (**Fig. S1A–C**). We focused on male WT and *Zfp687^−/−^* mice to specifically assess the effects of complete *Zfp687* loss on bone biology while minimizing potential confounding effects of sex hormones. At birth, *Zfp687^−/−^*mice appeared normal, with no evident macroscopic differences in body weight or length compared with WT littermates **(Fig. 1A)**. However, during postnatal development, *Zfp687^−/−^* mice showed an evident reduction in body size compared with WT controls. Based on this observation, we monitored body weight and length weekly from the first postnatal week in WT and *Zfp687^−/−^*mice (**Fig. 1B**). These analyses confirmed that complete *Zfp687* loss significantly reduced both parameters during postnatal growth. At 3 months of age, following the major phase of postnatal skeletal growth and medullary cavity expansion^34,35^, *Zfp687^−/−^* mice showed significant reductions in total body weight, body length, and femur and tibia lengths compared with WT littermates (**Fig. 1C-D**). However, organ weight-to-body weight ratios, including those of the liver and spleen, were comparable between genotypes indicating proportionate visceral organ growth (**Fig. S2A**).

**Figure 1:**
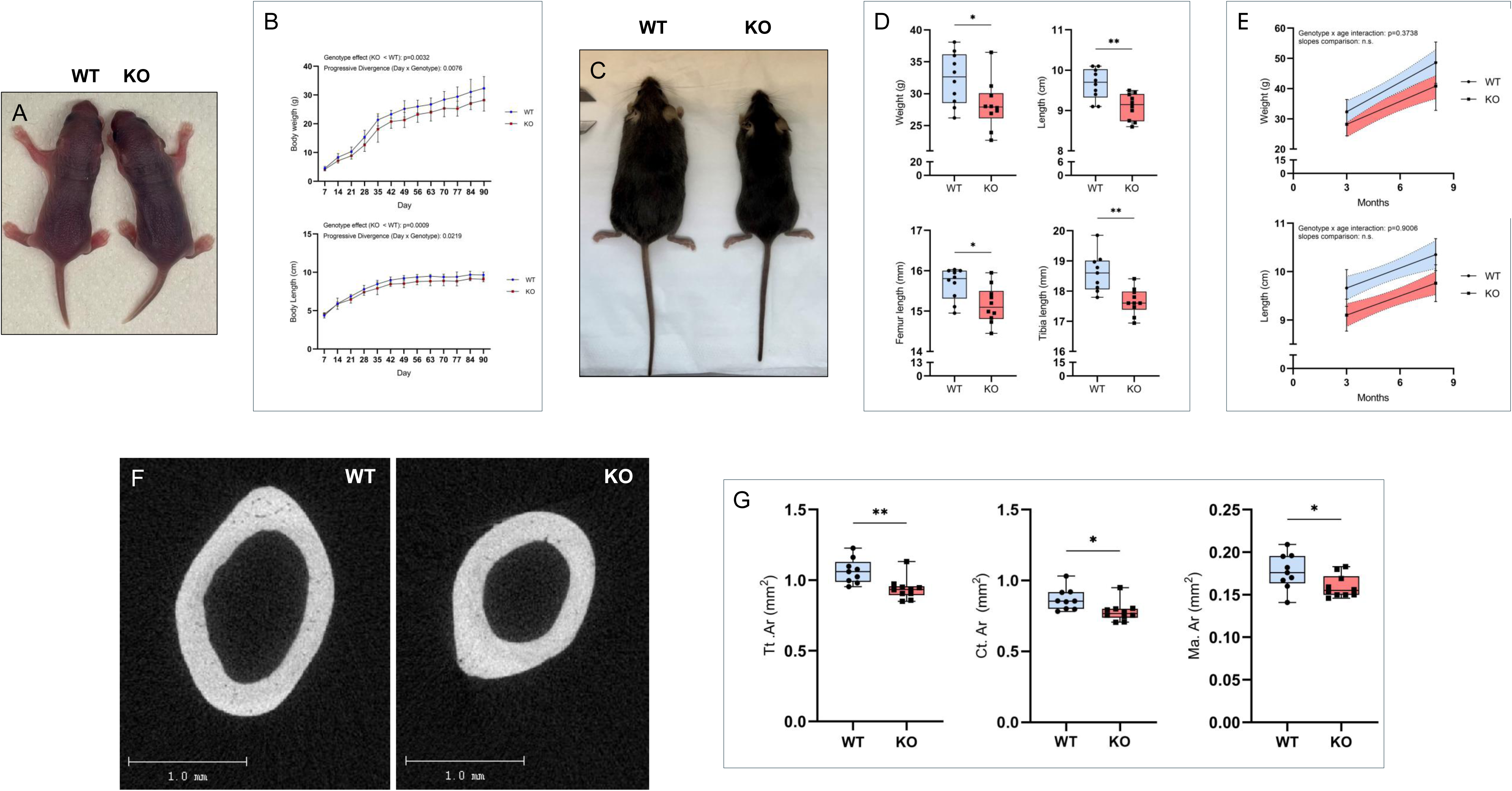
**A)** Representative image of 3-day postnatal WT and KO mice. **B)** Growth curves of body weight and length from postnatal day 7 to 90 of WT (n=10; blue dots) and KO (n=10; red squares) mice. Data are shown as mean±s.d. Body weight and length were analysed by fitting a mixed-effect model (REML). **C)** Representative image of 3-month-old KO and WT mice. **D)** Upper lane: body weight and length of 3-month-old WT and KO (n=10) mice. Lower lane: box plots showing the femur and tibia lengths of 3-month-old WT and KO (n=10) mice. Data are shown as mean±s.d. Statistical significance was assessed by two-tailed unpaired t test (*p<0.05; **p<0.01). **E-F)** Linear regression analysis of proportional growth in WT and KO mice between 3 and 8 months of age. Time (months) is shown on the x-axis, while body weight (**E**) and body length (**F**) on the y-axis. Independent cohorts were analysed at each time point (n=10 per genotype at 3 months of age; n=7 at 8 months of age). Simple linear regression was performed separately for WT and KO mice, and comparison of the regression slopes was no significantly different between genotypes. Shaded areas represent SD. Two-way ANOVA (genotype × age) revealed no significant interaction effect (p=03738; p=0.9006, respectively). **G)** Representative µCT cross-sectional images of femoral midshafts from 3-month-old WT and KO mice. Scale bar: 1 mm. **H)** Box plots showing the total cross-sectional area (Tt.Ar), cortical area (Ct.Ar), and medullary area (Ma. Ar) quantification at femoral midshafts from 3-month-old WT (n=9) and KO (n=10). Data are presented as the median±s.d. Statistical significance was assessed by two-tailed unpaired t test (*p<0.05; **p<0.01).

We confirmed the same phenotype at 8 months of age. *Zfp687*^−/−^ mice exhibited reduced body weight and body length compared with WT littermates, whereas internal organ weights remained proportional to body size (**Fig. S2B)**. We therefore examined longitudinal changes in body weight and length between 3 and 8 months. Both parameters increased progressively over time in WT and *Zfp687*^−/−^ mice, with parallel growth trajectories and no significant genotype × age interaction for either body weight (p=0.3738) or body length (p=0.9006) **(Fig. 1E)**.

Altogether, these data indicate that *ZNF687* is required for normal early postnatal somatic and skeletal growth. The absence of a genotype × age interaction between 3 and 8 months further suggests that the growth deficit is established during early postnatal development and subsequently maintained into adulthood, rather than progressively worsening with age. However, we observed significant differences in cortical geometry of long bones, consistent with the overall reduction in body size of *Zfp687*^−/−^ mice. Specifically, total cross-sectional area (Tt.Ar; p=0.0045), cortical area (Ct.Ar; p=0.0178), and marrow area (Ma.Ar; p=0.0394) were all significantly reduced in 3-month-old *Zfp687*^−/−^ mice compared with WT controls **(Fig. 1F–G)**, reflecting a smaller medullary cavity without detectable changes in overall bone mass at this stage. Consistently, micro-computed tomography (μCT) analysis of femurs from 3-month-old WT and *Zfp687*^−/−^ mice revealed no significant differences in trabecular or cortical bone mass parameters between genotypes **(Fig. S2C-D)**.

Given the established role of ZNF687 in bone remodelling, as demonstrated by our previous phenotypic characterization of the *Zfp687*^P937R^ knock-in mouse model, in which early alterations in osteoclast activity preceded age-dependent bone loss^20^, we next examined whether Zfp687 deficiency also resulted in dysregulated bone remodelling at 8 months of age. Quantitative µCT analysis of femurs from 8-month-old mice revealed a 50% increase in bone volume/total volume (BV/TV) in *Zfp687*^−/−^ mice compared with WT controls (p=0.0059) **(Fig. 2A–B; Fig. S3A)**. This increase was accompanied by higher trabecular thickness (Tb.Th; p=0.048) and trabecular number (Tb.N; p=0.0474), together with reduced trabecular separation (Tb.Sp; p=0.0497), indicating that trabecular bone accumulation in *Zfp687*^−/−^ mice was associated with broad changes in trabecular microarchitecture **(Fig. S3B)**. In contrast, cortical thickness (Ct.Th) was unchanged between genotypes at 8 months of age, indicating that the bone mass phenotype of Zfp687^−/−^ mice was restricted to the trabecular compartment **(Fig. S3B)**.

**Figure 2:**
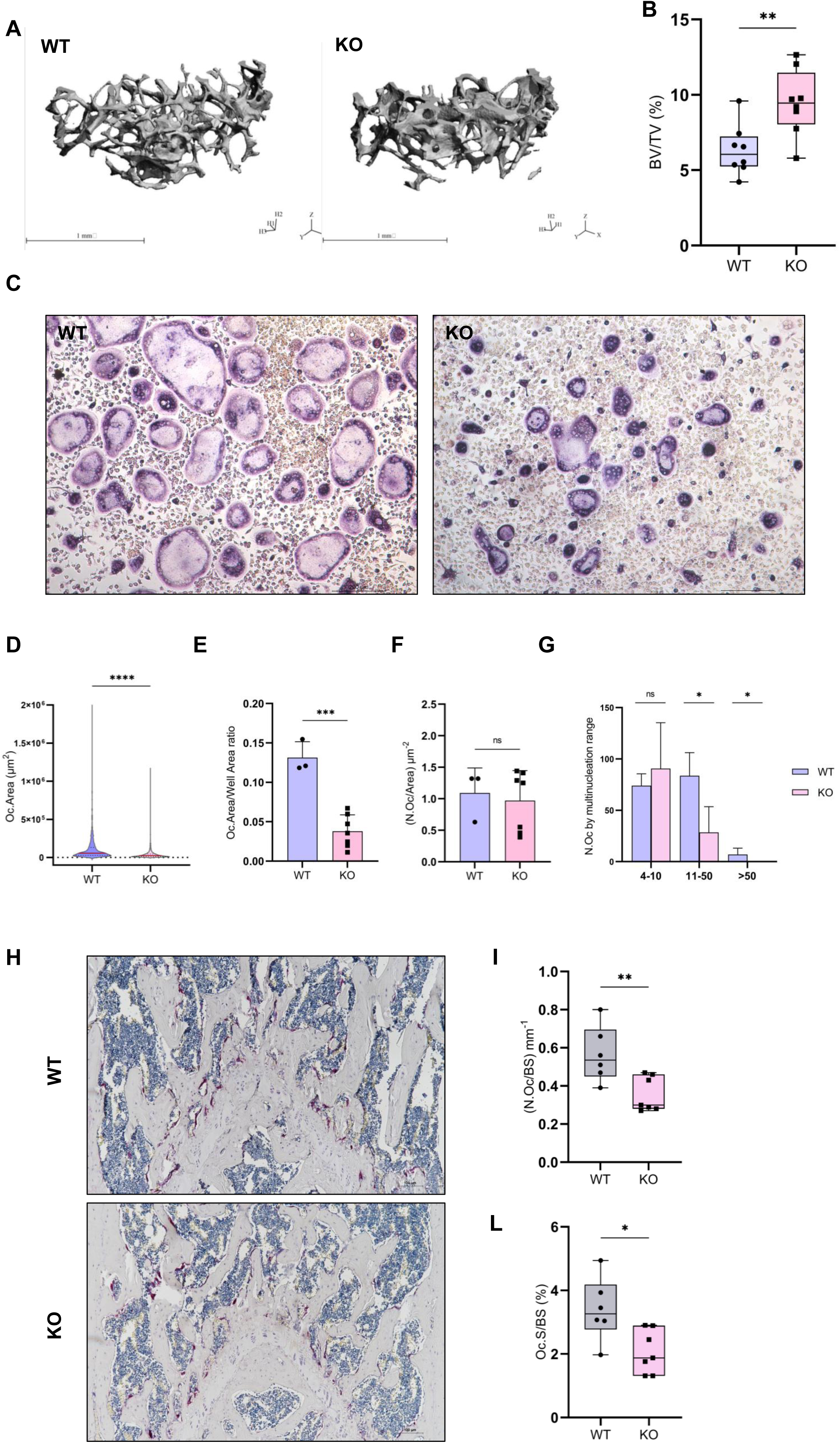
**A)** Representative µCT 3D reconstruction showing the trabecular bone in the distal femoral epiphysis of 8-month-old WT and KO mice. Scale bars: 1 mm. **B)** Box plots showing the BV/TV percentage measured by µCT in the trabecular bone of femoral distal epiphysis of WT and KO (n=8) mice. Data are presented as median±s.d. and statistical significance was assessed by two-tailed unpaired t-test (**p<0.01). **C)** Representative TRAP^+^ OCs from WT and KO BMMs differentiation. Scale bars: 200 µm. **D)** Violin plots with median (central red line), first (lower dotted black line), and third quartiles (upper dotted black line) of differentiated TRAP^+^ OCs area derived from WT (n=3) and KO (n=7). Data represent area measurements from n=1002 WT and n=1749 KO OCs. Statistical significance was assessed by two-tailed unpaired t-test (****p<0.0001). **E)** Bar graph showing TRAP^+^ OC-covered surface WT and KO cells, quantified as the sum of individual TRAP^+^ cell areas normalized to the total analysed field area. Data are shown as mean±s.d. Statistical significance was assessed by two-tailed unpaired t test (*p<0.05). **F)** Bar graph showing OC density expressed as OC number normalized to the analysed field area. Data are shown as mean±s.d. Statistical significance was assessed by two-tailed unpaired t test (ns=not significative). **G)** Bar graph showing OC stratified by nuclear number (4-10, 11-50, and >50 nuclei) from WT and KO. Data are shown as mean±s.d. Statistical significance was assessed by multiple unpaired t test followed by the Holm-Šídák method for correction (ns=not significative; *adj. p<0.05). **H)** Representative TRAP^+^-stained sections of femoral distal epiphysis from 3-month-old WT and KO mice showing TRAP^+^ OCs in purple; nuclei were counterstained with hematoxylin. Scale bars: 100 µm. **I)** Box plots showing the OC surface to bone surface ratio (Oc.S/BS), and **L)** the OC number to bone surface per mm ratio (Oc.N/BS) in 3-month-old WT and KO (n=7) mice. Data are presented as the median±s.d. Statistical significance was assessed by two-tailed unpaired t test (*p<0.05; **p<0.01).

To determine whether impaired osteoclast differentiation underlies this age-dependent bone phenotype, we next assessed osteoclastogenesis *in vivo* and *in vitro* at 3 months of age. We isolated bone marrow cells and expanded primary bone marrow monocytes/macrophages (BMMs) in the presence of M-CSF. Osteoclast differentiation was then induced by stimulation with murine RANKL for 5 days. *Zfp687*^−/−^ BMMs markedly showed a reduced capacity to undergo osteoclast differentiation. Quantification of TRAP^+^ multinucleated cells containing ≥4 nuclei showed that *Zfp687*^−/−^ osteoclasts were significantly smaller (p<0.0001) and covered a reduced surface area (p=0.0002) compared with WT controls **(Fig. 2C–E)**. However, despite the smaller area, *Zfp687*^−/−^-derived cultures showed same osteoclast density per area as WT cells (**Fig. 2 F**). Indeed, stratification of TRAP^+^ multinucleated cells by nuclei number revealed that the formation of small osteoclasts (4–10 nuclei) was preserved in *Zfp687*^−/−^ cultures. However, Zfp687 deficiency led to a significant reduction in mature osteoclasts containing 11–50 nuclei (p=0.0446). Most strikingly, the late cell-fusion required for the formation of giant osteoclasts (>50 nuclei) was completely abolished in *Zfp687*^−/−^ cells (p=0.0446) **(Fig. 2G)**. Together, these data demonstrate that *Zfp687* deficiency severely impairs overall osteoclastogenesis and full maturation into multinucleated and giant osteoclasts. In line with these results, *Zfp687* loss strongly reduced osteoclastogenesis *in vivo*, as evidenced by histological evaluation of TRAP^+^ osteoclasts in *Zfp687*^−/−^ bone sections compared with WT controls (**Fig. 2H**). Histomorphometric analysis confirmed these observations: osteoclast surface per bone surface (Oc.S/BS) but also osteoclast number per bone surface (N.Oc/BS) were reduced by 39% (p=0.0158) and 37% (p=0.0096) respectively, in femoral sections from *Zfp687*^−/−^ mice compared with WT animals (**Fig. 2I-L**). Since equal number of precursors were seeded in vitro, these in vivo data suggest that Zfp687 loss affects not only osteoclast differentiation, but also the pool of cells committed toward the osteoclastogenic lineage. This phenotype mirrors, in the opposite direction, the enhanced early osteoclastogenesis and subsequent age-dependent bone loss previously observed in the *Zfp687*^P937R^ knock-in model. These complementary gain- and loss-of-function data identify ZNF687 as a critical regulator of osteoclastogenesis in adulthood, with effects on age-dependent bone remodelling. Additionally, these results suggest that ZNF687-dependent osteoclast activity contributes to early postnatal skeletal development, when osteoclast-mediated expansion of the medullary cavity is required for proper BM maturation.

### Zfp687 controls bone marrow myeloid populations with osteoclastogenic potential

Several metabolic bone disorders, including PDB, are associated with dysfunctional osteoclast activity and abnormal bone remodelling^17,36,37^. To assess whether impaired osteoclastogenesis in *Zfp687*^−/−^ mice reflected changes in the osteoclast precursor pool, we analysed the BM cellular compartment of 3-month-old mice, which corresponds to the stage of reduced *in vivo* osteoclast formation. Total BM cellularity was significantly reduced in *Zfp687*^−/−^ mice compared with WT littermates (p=0.0133), consistent with the reduced femoral marrow area observed in these mice **(Fig. 3A)**. Importantly, this reduction remained substantial (by 23%) after normalization to bone length, indicating that the decreased cellularity reflects an alteration of the BM cellular compartment upon *Zfp687* loss, rather than a consequence of reduced skeletal size.

**Figure 3:**
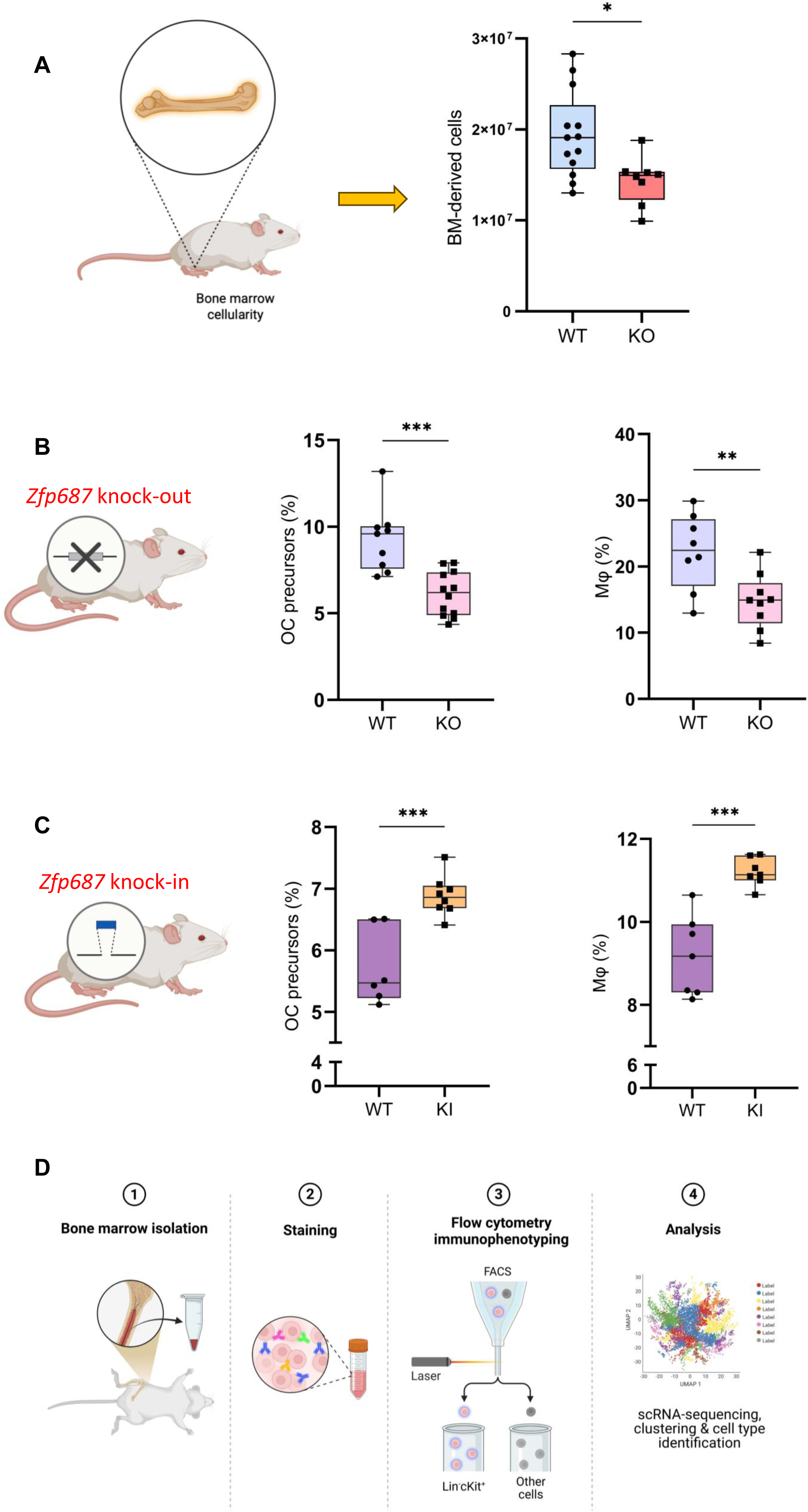
**A)** Box plots showing the total BM cell yielded from 3-month-old WT (n=13) and KO (n=8) mice. Statistical significance was assessed by two-tailed unpaired t test (*p<0.05). **B)** Box plots showing BM-derived OC precursors and F4/80^+^ Mφ frequencies in 3-month-old WT (n=9) and KO (n=12) mice. Data are presented as the median±s.d. Statistical significance was assessed by two-tailed unpaired t test (**p<0.01; ***p<0.001). **C)** Box plots showing BM-derived OC precursors and F4/80^+^ Mφ frequencies in 3-month-old WT (n=6) and KI (n=8) mice. Data are presented as the median±s.d. Statistical significance was assessed by two-tailed unpaired t test (***p<0.001). **D)** Schematic workflow illustrating BM isolation, Ter119^−^B220^−^CD117^+^ cell sorting by FACS, and scRNA-sequencing library preparation.

Since osteoclasts derive from BM-resident myeloid progenitors, we next evaluated the impact of *Zfp687* deficiency on osteoclast precursors using a flow cytometric approach. Common monocyte progenitors (cMoP) have been described as major source of osteoclasts both *in vitro* and *in vivo*^6,8–10,38^. Therefore, we evaluated the frequency of osteoclast progenitors by their Ter119^−^B220^−^CD117^+^CD115^+^CD11b^−^ immunophenotype. We found a significant reduction affecting osteoclast precursor cells in *Zfp687*^−/−^ BM compared to WT (p=0.0002) at 3 months of age (**Fig. 3B, Fig. S4A**). Given that osteoclast precursor cells can also give rise to macrophages (Mφ), which retain osteoclastogenic potential upon M-CSF and RANKL stimulation^39,40^, we further assessed F4/80^+^ Mφ abundance. This analysis also revealed a significant reduction of F4/80^+^ Mφ in BM from *Zfp687*^−/−^mice compared to WT littermates (p=0.0074; **Fig. 3B, Fig. S4B**). Together, these results indicate that the reduced number and formation of osteoclasts observed in *Zfp687*^−/−^ bone sections are, at least in part, driven by a reduced availability of BM myeloid precursors with osteoclastogenic potential, identifying a previously unrecognized role for ZNF687 in modulating BM myeloid lineage commitment.

To directly compare the effects of *Zfp687* deficiency and the pagetic P937R mutation on BM myeloid populations, we applied the same flow cytometric strategy to 3-month-old *Zfp687*^P937R^-KI mice. In striking contrast to the *Zfp687*-KO model, we observed that the P937R mutation promoted a marked expansion of both osteoclast precursors and F4/80^+^ Mφ compared to WT controls (p=0.0007; p=0.0002, respectively) **(Fig. 3C, Fig. S4C-D)**. This expansion is consistent with the enhanced *in vivo* osteoclastogenesis observed at 3 months of age in the KI mouse model, which precedes the bone loss phenotype we previously reported^20^, and mirrors, in an opposite direction, the osteoclastogenic progenitors deficit seen in *Zfp687^−^*^/−^ mice.

Taken together, these findings establish a relationship between ZNF687 and the frequency of BM-resident populations with osteoclastogenic potential, with important implications for the early pathogenic events driving the severe ZNF687-related PDB.

### Zfp687 loss selectively disrupts the early myeloid progenitor compartment

To determine whether Zfp687 loss altered the broader BM myeloid progenitor compartment, we performed single-cell RNA sequencing (scRNA-seq) on sorted BM-derived Ter119^−^B220^−^CD117^+^ cells from 2 *Zfp687*^−/−^ and 2 WT mice **(Fig. 3D, S4E)**. The single-cell transcriptomic profile was performed on 30600 high-quality singlets, obtained after quality control filtering, doublet removal, SCTransform normalization and cell cycle phase distribution (**Fig. S5A-C)**. Unsupervised clustering identified 22 transcriptionally distinct clusters, spanning the major myeloid lineages expected from this sorting strategy: HSC/MPP (multipotent progenitors), granulocyte/monocyte progenitors (GMP) and more mature cells, as well as a minor erythroid progenitor population **(Fig. 4A)**. Cell type annotation was performed by integrating canonical lineage marker expression with label transfer from the Lin^−^Kit^+^ murine BM myeloid reference dataset deposited by Paul and colleagues^41^ (**Fig. S5D**). The annotated clusters recapitulated the known myeloid differentiation hierarchy, with HSC/MPP at the apex, followed by cycling GMP and promyelocyte populations, and more terminally differentiated neutrophils and monocytes (**Fig. 4B**). Comparative UMAP and stacked bars composition analysis revealed a striking reduction of cycling myeloid progenitor populations in *Zfp687*^−/−^ samples compared to WT, most prominently within the GMP compartment (**Fig. 4A, C**). To quantify these composition differences, we performed differential abundance analysis using propeller, which models cluster proportions among biological replicates accounting for inter-sample variability. Interestingly, we found that one of the most significantly depleted clusters (FDR<0.05) upon *Zfp687* loss was the early GMP cycling cluster (cluster 4, FDR=0.038, PropRatio=9.92), which was reduced approximately 10-fold relative to WT (**Fig. 4D)**. This finding is of biological relevance, as GMPs represent the common myeloid progenitor from which both the granulocytic and monocyte/macrophage lineages diverge, the latter giving rise to osteoclast precursors in BM^15,42,43^. Congruently, the second cluster significantly reduced in KO-derived BM was the Promyelocyte/myelocyte transition cluster (cluster 3, FDR=0.038, PropRatio=2.29), which directly derives from the early GMP cycling cluster. Additionally, no clusters were significantly enriched in *Zfp687*^−/−^ mice compared to WT, indicating the absence of compensatory mechanisms maintaining BM cellularity after *Zfp687* depletion. Thus, the impressive reduction of GMPs pool in *Zfp687*^−/−^ mice provides a mechanistic basis for the impaired macrophage and osteoclast precursor abundance previously observed by flow cytometry.

**Figure 4:**
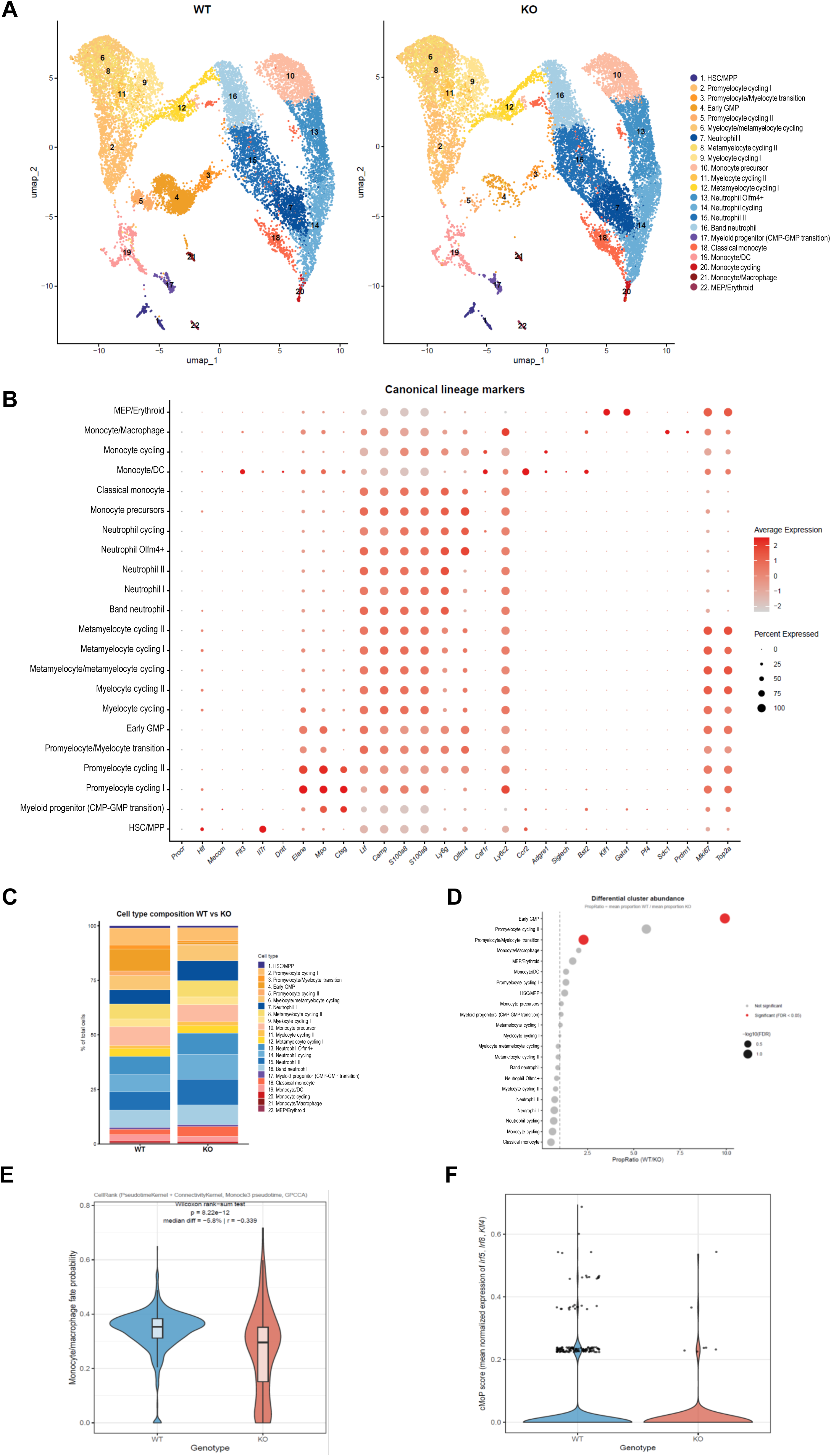
**A)** Split UMAP visualization comparing cell type distribution between WT and KO mice. Cells are colored by annotated cell type identity across 22 transcriptionally distinct clusters. **B)** DotPlot showing scaled average expression (color) and percentage of expressing cells (dot size) for canonical lineage markers across the 22 distinct clusters. **C)** Stacked bar plot showing cell type composition as percentage of total cells, comparing WT and KO mice. **D)** Differential abundance analysis of the 22 clusters between WT and KO mice (PropRatio=mean proportion WT/KO; red dots indicate significantly altered clusters with FDR<0.05). **E)** Violin plot showing monocyte/Mφ fate probability inferred by CellRank in the early GMP cluster of WT and KO mice. Statistical significance was assessed by Wilcoxon rank-sum test; effect size as rank-biserial correlation (r). **F)** Violin plot showing the cMoP transcriptional score (mean normalized expression of *Irf5*, *Irf8*, *Klf4*) in the early GMP cluster of WT and KO mice. Points indicate cells with score >0.05.

To determine whether the reduction in GMPs could reflect a block toward myeloid differentiation, we performed trajectory inference using Monocle3, rooting the analysis at the HSC/MPP cluster. The learned trajectory recapitulated the expected myeloid differentiation continuum, with pseudotime values increasing progressively from HSC/MPP through cycling progenitors to mature neutrophils and monocytes (**Fig. S5E-F**). Notably, the trajectory topology and branch point structure were preserved between genotypes. However, to explore whether *Zfp687* deficiency affects the inferred lineage bias of progenitor cells, we inferred cellular fate probability toward the monocyte/macrophage lineage using CellRank 2^44^. Notably, fate probability analysis across the GMP cluster disclosed a significant reduction in monocyte/macrophage fate probability in *Zfp687*^−/−^ cells (p=8.22e^-12^; r=-0.339; **Fig. 4E**), in line with the lower production of osteoclast precursors and macrophages. Consistently, based on a published cMoP transcriptional signature (*Irf5*, *Irf8*, *Klf4*)^42^, the fraction of cMoP marker expressing cells within the GMP compartment was reduced in *Zfp687*^−/−^ mice, compared to WT (**Fig. 4F**).

Collectively, these findings indicate that ZNF687 is required for the proper maintenance of myeloid progenitors in BM, with the GMP compartment being the most severely affected. The compromised myeloid lineage compartment suggests that the altered osteoclast precursor and macrophage frequencies previously observed may originate, at least in part, from a disruption at the level of myeloid-committed progenitors.

### Zfp687 loss impairs BRD4 signalling and M-CSF-dependent osteoclastogenesis

Given the role of bromodomain-containing proteins in myeloid lineage commitment and osteoclastogenesis^27,28^, we investigated whether ZNF687-dependent alterations were associated with changes in BRD4 and downstream osteoclastogenic regulators at the protein level. To this end, we analysed BRD4 levels in *Zfp687*^+/−^ RAW264.7 cells, which we previously showed to be unable to undergo efficient osteoclast differentiation upon RANKL stimulation^20^. We observed that *Zfp687*^+/−^clones exhibited a consistent reduction in BRD4 protein levels compared with WT cells by western blot analysis **(Fig. 5A–B)**. This was accompanied by a significant reduction in c-Myc, a well-established transcriptional target of Brd4^28,45–47^ and a key regulator of osteoclastogenesis through the modulation of *NFATc1* expression^26,28,48,49^, as well as a consistent, reduction in NFATc1 protein levels, a master transcription factor required for osteoclast commitment and differentiation^50^ (**Fig. 5A**). To validate these observations in a primary cell context, we analysed BRD4, c-Myc, and NFATc1 protein levels in BMMs derived from *Zfp687*-KO mice. Consistently, BRD4 protein levels were significantly reduced in *Zfp687*^−/−^-derived BMMs compared to WT controls (p=0.0448), accompanied by marked reduction in c-Myc (p=0.0034), and NFATc1 (p=0.0099) levels (**Fig. 5B**).

**Figure 5:**
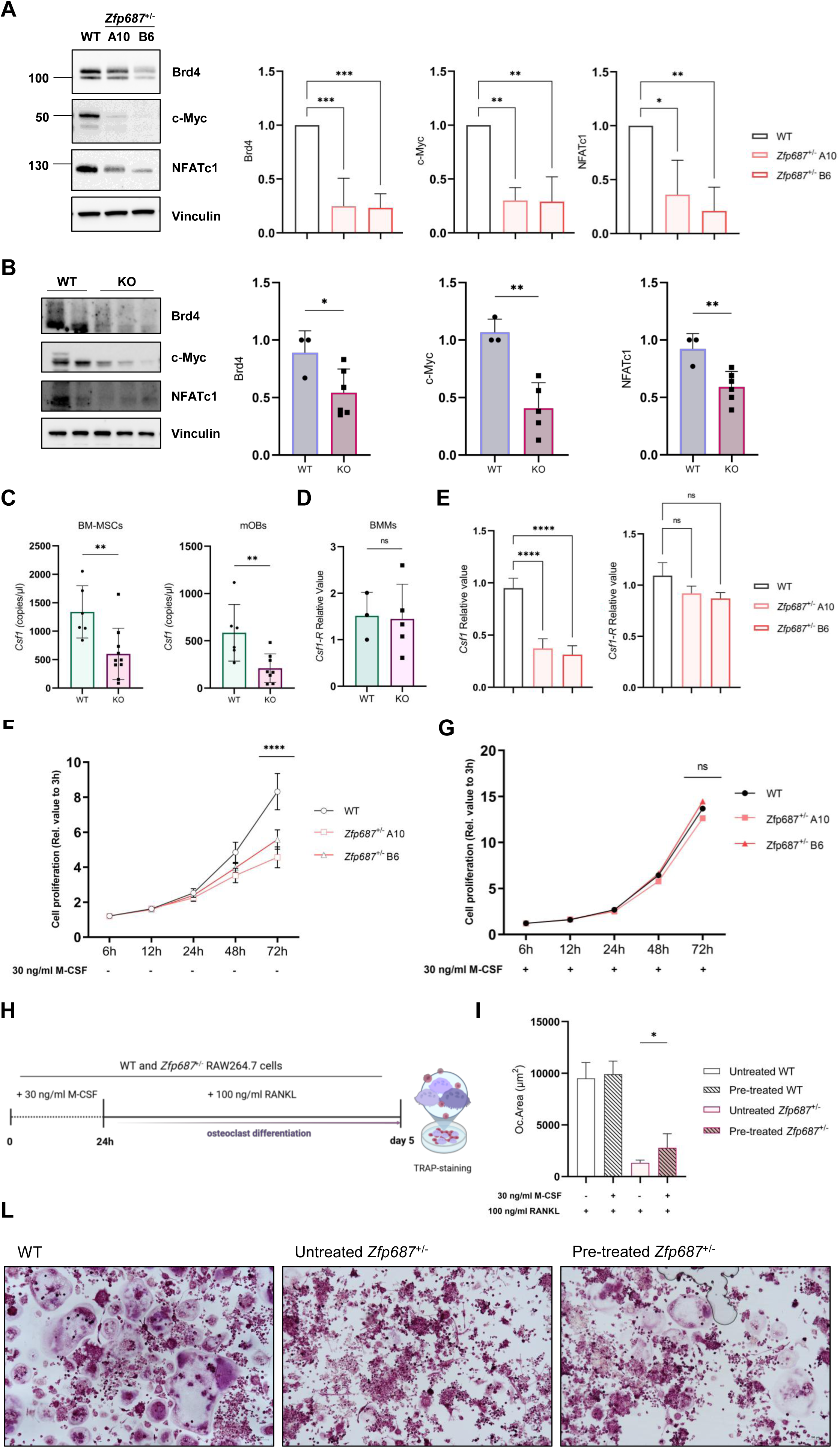
**A)** Representative western blot analysis of Brd4, c-Myc, and NFATc1 protein levels in WT and *Zfp687*^+/−^ RAW264.7 clones (A10 and B6). Vinculin was used as loading control. Relative quantification of Brd4, c-Myc, and NFATc1 protein levels. Data represent the mean±s.d. Statistical significance was assessed by one-way ANOVA with Dunnett’s multiple comparisons test vs WT (*p<0.05; **p<0.01; ***p<0.001; ns=not significative). **B)** Representative western blot analysis of Brd4, c-Myc, and NFATc1 protein levels in WT and KO BMMs. Relative quantification of Brd4, c-Myc, and NFATc1 protein levels. Data represent the mean±s.d. Statistical significance was assessed by two-tailed unpaired t test (*p<0.05; **p<0.01). **C)** Bar graphs showing ddPCR results expressed in copies/µl of *Csf1* expression in BM-MSCs (left) and mOBs (right) from WT (n=6) and KO (n=8;10) mice. Data are presented as mean±s.d of two independent experiments. Statistical significance was assessed by two-tailed unpaired t test (**p<0.01). **D)** Bar graphs showing qRT-PCR results of *Csf1-R* expression in WT (n=3) and KO (n=5) derived BMMs. Data are presented as mean±s.d of two independent experiments. Statistical significance was assessed by two-tailed unpaired t test (ns=not significative). **E)** Bar graphs showing qRT-PCR results of *Csf1* expression (left) and *Csf1-R* expression (right) in WT and *Zfp687*^+/−^ RAW264.7 clones. Data are presented as mean±s.d of 2 independent RNA extractions. For each extraction, qRT-PCR was performed in 6 technical replicates. Statistical significance was assessed by one-way ANOVA with Dunnett’s multiple comparisons test vs WT (ns=not significative; *p<0.05; ****p<0.0001). **F)** CCK8 proliferation of WT and *Zfp687*^+/−^ RAW264.7 clones at 6, 12, 24, 48, and 72 hours from the start of the assay (3h, relative time) ± 30 ng/ml of M-CSF. Data are shown as the mean±s.d of 4 independent experiments. Statistical significance was assessed by two-way ANOVA (****p<0.0001; ns= not significative). **G)** Schematic representation of the experimental design. **H)** Quantification of TRAP⁺ multinucleated OC area. Bar graphs showing WT and the mean of *Zfp687*^+/−^ RAW264.7 clones pretreated ± 30 ng/ml M-CSF for 24 hours. Data are presented as mean±s.d. from 2 independent experiments. Statistical analysis was performed using an unpaired two-tailed Student’s t-test (*p<0.05). WT cells served as control of OC differentiation. **I)** Representative images of OC differentiation preceded by ± 30 ng/ml M-CSF for 24 hours. Images show WT cells (left), untreated (middle) and M-CSF-pretreated *Zfp687*^+/−^ cells (right). Scale bar, 100 µm.

We next investigated whether Zfp687 deficiency also affects extrinsic niche-derived signals that support maintenance of the OC progenitor pool, including the GMP compartment, by analysing M-CSF, a critical regulator of myeloid cell survival, proliferation, and differentiation^42,51–57^. We therefore isolated and expanded BM cells to obtain BM mesenchymal stromal cells (BM-MSCs), which were subsequently induced toward osteoblastic differentiation. Digital droplet PCR (ddPCR) analysis revealed a significant reduction in *Csf1* expression in both BM-MSCs (p=0.007) and mOBs (p=0.0094) from *Zfp687*^−/−^ mice compared with WT controls **(Fig. 5C)**, while *Csf1r* expression was unchanged in BMMs derived from *Zfp687*^−/−^ mice compared with WT controls **(Fig. 5D)**. These results suggest that reduced M-CSF expression in Zfp687-deficient stromal and osteoblastic cells may link the BM microenvironment to the reduced OC precursor and F4/80^+^ macrophage populations observed *in vivo*.

To determine whether reduced M-CSF availability contributes to the osteoclastogenic defects associated with *Zfp687* deficiency, we used *Zfp687*^+/−^ RAW264.7 cells as a cell-autonomous model that expresses basal levels of M-CSF and relies on an autocrine feedback loop for proliferation and partial differentiation^58,59^. Gene expression analysis showed a marked reduction in *Csf1* levels in *Zfp687*^+/−^ clones compared with WT cells, whereas expression of its receptor, *Csf1-R*, was unaffected **(Fig. 5E)**. Given the pivotal role of M-CSF in promoting myeloid cell survival and proliferation, we next evaluated the proliferative capacity of *Zfp687*^+/−^ RAW264.7 cells. These cells exhibited significantly reduced proliferation compared with WT cells, as assessed by CCK8 assay at multiple time points (6, 12, 24, 48, and 72 h) **(Fig. 5F)**. Notably, supplementation with exogenous M-CSF fully rescued the proliferative deficit of *Zfp687*^+/−^ RAW264.7 cells to levels comparable with WT cells **(Fig. 5G)**. We then asked whether exogenous M-CSF could also ameliorate the osteoclast differentiation defect in *Zfp687*^+/−^ RAW264.7 cells. Pretreatment of *Zfp687*^+/−^ cells with exogenous M-CSF prior to RANKL stimulation improved OC formation compared with untreated *Zfp687*^+/−^ cells (p=0.0457) **(Fig. 5H-L)**, although differentiation was not fully restored to WT levels. The same pretreatment had no effect on WT osteoclastogenesis.

Together, these data support a role for ZNF687 in regulating M-CSF expression both in a cell-autonomous context and within the BM microenvironment, with functional consequences for myeloid cell proliferation and OC differentiation. These findings are consistent with the reduced BM myeloid populations and impaired osteoclastogenesis observed in the Zfp687-KO mouse model.

### The pagetic P937R mutation enhances clonogenic haematopoietic output in human progenitors

Based on our findings that *Zfp687* mutant mice exhibit marked alterations within BM myeloid populations, we sought to determine whether human haematopoietic progenitors carrying the pagetic P937R mutation display altered clonogenic output and lineage potential.

To address this, we generated human induced pluripotent stem cells (hiPSCs) from dermal fibroblasts derived from a P937R-mutant pagetic patient and from his unaffected brother, who served as matched sibling control (Ctrl). Representative ZNF687-P937R and Ctrl hiPSC clones displayed a normal karyotype and expressed the pluripotency markers NANOG, and OCT3/4 **(Fig. 6A)**. We then subjected P937R- and Ctrl-derived hiPSCs to haematopoietic differentiation, first inducing mesoderm specification and subsequently generating haematopoietic progenitor cells (HPCs) **(Fig. 6B)**. After 12 days of differentiation, FACS analysis of HPCs, identified as CD43^+^CD34^+^CD45^+^ cells, showed comparable frequencies between P937R mutant- and Ctrl-differentiated cells (**Fig. 6C**). We next subjected P937R mutant- and Ctrl-derived HPCs to colony-forming unit assays to assess erythroid progenitors, including CFU-E and BFU-E, granulocyte–macrophage progenitors (CFU-GM), and multipotent granulocyte–erythroid–macrophage–megakaryocyte progenitors (CFU-GEMM). P937R mutant-derived colonies emerged earlier and were more numerous than Ctrl colonies at day 9 of differentiation. By day 15, P937R mutant-derived HPCs maintained a higher clonogenic output, generating a 44% increase in total CFU-colonies compared with Ctrl-HPCs. In addition, P937R-derived colonies resulted markedly larger than Ctrl colonies, suggesting that the P937R mutation confers enhanced proliferative potential together with an expanded clonogenic progenitor pool **(Fig. 6D)**. Analysis of colony-type distribution revealed that the P937R mutation altered clonogenic kinetics and lineage distribution. At day 15, P937R mutant-derived HPCs showed a striking expansion of multipotent CFU-GEMM colonies (increasing from 8% to 26%, representing a 3-fold enrichment), alongside a 2-fold increase of BFU-E colonies. In contrast, the distribution frequency of more committed CFU-GM colonies was reduced up to 42% in P937R mutant-derived cells compared with Ctrl cells. However, despite their lower relative frequency, P937R mutant-derived CFU-GM colonies displayed a distinct phenotype, appearing larger and markedly more cellular than Ctrl CFU-GM colonies. Representative images showed that Ctrl CFU-GM formed modest clusters of large, distinguishable cells, whereas P937R mutant-derived CFU-GM generated dense aggregates of tightly packed cells, consistent with enhanced proliferative expansion.

**Figure 6:**
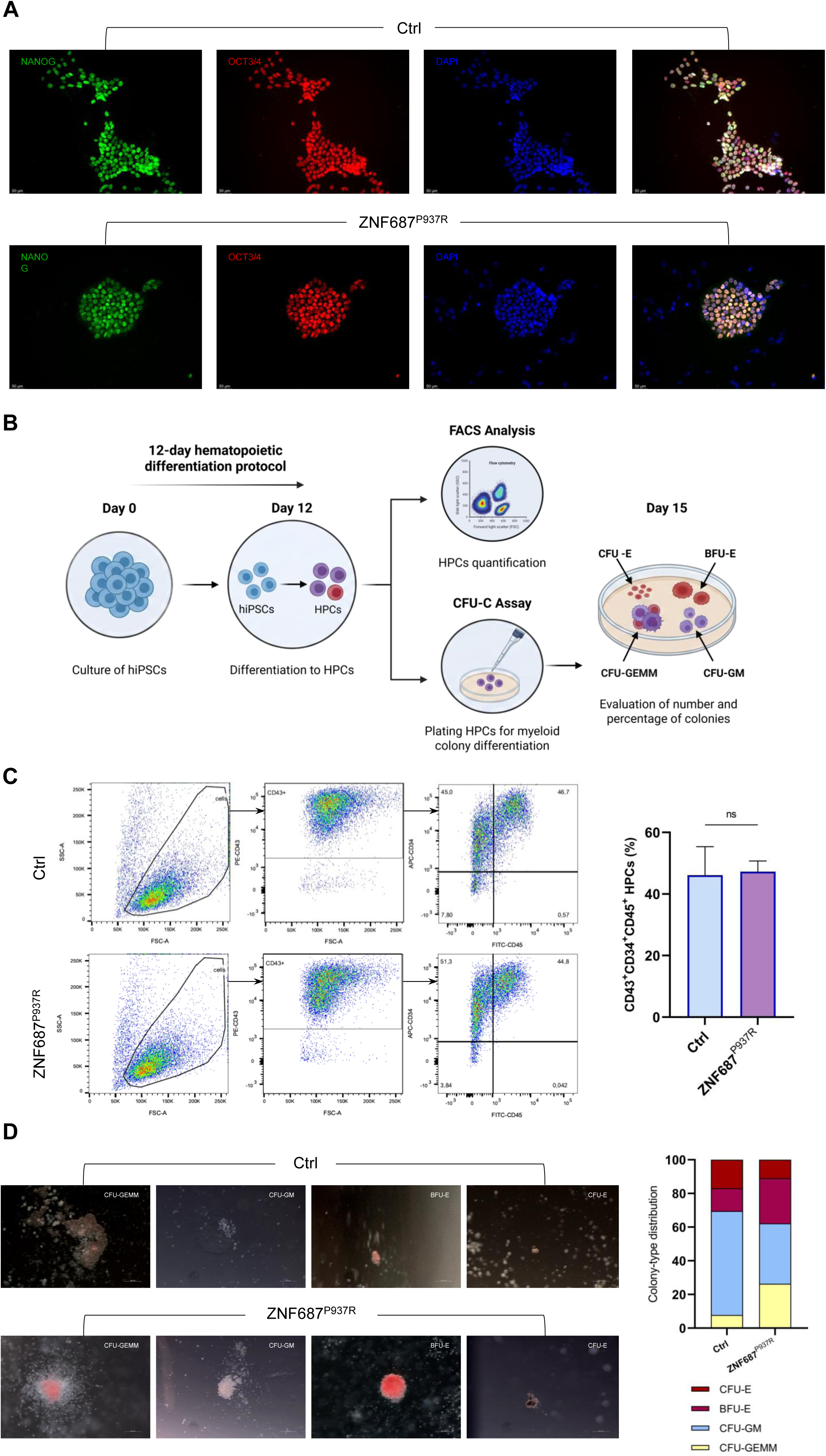
**A)** Expression of pluripotency markers NANOG and OCT3/4 measured by IF in the healthy-donor and P937R-mutant PDB patient derived clones. **B)** Schematic workflow for hiPSC hematopoietic differentiation and functional assessment of HPCs. hiPSCs were subjected to a 12-day hematopoietic differentiation to generate HPCs. At day 12, the percentage of HPCs was quantified by flow cytometry (top); HPCs were plated in methylcellulose medium and cultured for 15 days for a CFU-C assay (bottom). **C)** HPCs flow cytometry characterization at day 12 of differentiation. HPCs were gated on the CD43⁺ population, and CD34⁺CD45⁺ cells are reported as percentages. Representative contour plots are shown for the healthy donor clone (Ctrl, left) and the P937R-mutant PDB patient clone (ZNF687^P937R^, right). **D)** Representative images of colony-forming unit (CFU-C) assays (left). Stacked bar chart showing the frequence of colony-type distribution for the Ctrl- and the ZNF687^P937R^-derived clones. Colonies were classified as CFU-GEMM, CFU-GM, BFU-E and CFU-E, and colour-coded as indicated.

Thus, the skewed colony distribution, together with the increased size and cellularity of P937R-derived colonies, suggests that the pagetic mutation enhances clonogenic haematopoietic output and favours expansion of early multipotent progenitor states. These findings are consistent with the expanded BM myeloid populations observed in the Zfp687^P937R^ knock-in model and support a gain-of-function effect of the P937R mutation on the haematopoietic progenitor compartment.

## Discussion

In this study, we define ZNF687 as a regulator of BM myeloid progenitor dynamics that functionally couples haematopoietic homeostasis to osteoclastogenesis in PDB. Using complementary loss- and gain-of-function models, we show that *Zfp687* ablation reduces BM cellularity, depletes GMPs, osteoclast progenitors and macrophages, and impairs osteoclastogenic differentiation, ultimately resulting in increased trabecular bone mass during adulthood. Conversely, the opposite phenotype observed in the *Zfp687*^P937R^-KI model supports a dosage-sensitive role of ZNF687 in controlling the size and osteoclastogenic competence of the BM myeloid compartment. These findings extend the previously described role of ZNF687 in osteoclast differentiation^20^ and reposition this factor upstream of mature osteoclast formation, at the level of the progenitor populations that sustain pathological bone remodelling.

A central finding of this work is that *Zfp687* loss selectively affects the early myeloid progenitor compartment. Single-cell profiling of sorted BM-derived c-Kit^+^ haematopoietic progenitors revealed a significant depletion of the early cycling GMP cluster in *Zfp687*-KO mice, without evidence of a global block in myeloid differentiation. Fate-probability analysis further indicated reduced bias toward the monocyte/macrophage lineage, paralleled by contraction of a cMoP-primed subfraction. This single-cell prediction was supported by flow cytometry, which showed reduced osteoclast progenitors and F4/80^+^ macrophages in KO marrow. Thus, *Zfp687* loss affects the osteoclastogenic myeloid compartment at two levels: quantitatively, by reducing progenitor abundance, and qualitatively, by impairing their capacity to progress toward efficient osteoclast differentiation. The expansion of osteoclast progenitors and macrophages in *Zfp687*^P937R^-KI mice reinforces this model and suggests that the pagetic mutation acts, at least in part, by expanding the myeloid reservoir from which osteoclasts arise.

The human iPSC-derived haematopoietic differentiation data further support this concept. P937R-derived progenitors showed accelerated colony emergence, increased clonogenic output, enrichment of primitive/multipotent CFU-GEMM colonies, and enlarged hypercellular myeloid colonies. Although these data do not indicate a simple increase in terminal myeloid commitment, they support a gain-of-function effect of the P937R mutation on the clonogenic and proliferative behaviour of haematopoietic progenitors. Together with the *in vivo* KI data, these findings suggest that ZNF687-related PDB may not originate solely from hyperactive mature osteoclasts, but also from earlier alterations in haematopoietic progenitor output and lineage potential. This provides a mechanistic framework for the aggressive osteoclastogenic phenotype associated with the P937R mutation.

At the molecular level, *Zfp687*-deficient cells showed reduced BRD4 protein levels, accompanied by decreased c-Myc and NFATc1 expression. BRD4 and c-Myc are established regulators of the osteoclastogenic transcriptional programme, particularly through control of NFATc1 induction^26,28,48,60,61^. However, while previous studies have mainly positioned this axis downstream of RANKL stimulation, our data suggest that ZNF687 modulates BRD4–c-Myc–NFATc1 signalling at baseline, before osteoclastogenic induction. This implies that ZNF687 may contribute to establishing the basal transcriptional competence of osteoclast precursors, thereby determining their ability to mount a full NFATc1-driven differentiation response upon RANKL exposure. In this model, reduced BRD4, c-Myc and NFATc1 levels in *Zfp687*-deficient cells would generate a less permissive progenitor state, consistent with the impaired osteoclastogenesis observed in both *Zfp687*^+/−^RAW264.7 cells and primary *Zfp687*^−/−^ BMMs.

In parallel, our data support a niche-dependent mechanism through which ZNF687 regulates BM myeloid progenitor availability. *Csf1* expression was reduced in BM-MSCs and osteoblasts from *Zfp687*-KO mice, as well as in *Zfp687*^+/−^ RAW264.7 cells, whereas *Csf1-R* expression remained unchanged. Since M-CSF is essential for the survival, proliferation and differentiation of myeloid progenitors and osteoclast precursors^42,51–57^, reduced stromal M-CSF production likely contributes to the contraction of the osteoclastogenic compartment observed *in vivo*. Functionally, exogenous M-CSF fully rescued the proliferative defect of *Zfp687*^+/−^ RAW264.7 cells, but only partially restored osteoclast formation upon RANKL stimulation. This incomplete rescue supports a dual model in which ZNF687 controls osteoclastogenesis through both a stromal/niche-derived M-CSF signal that sustains precursor availability and a cell-autonomous BRD4–c-Myc–NFATc1 axis that determines differentiation competence. However, the molecular link between ZNF687 and BRD4 remains to be defined, including whether ZNF687 regulates BRD4 transcription, protein stability, or acts indirectly through intermediate effectors. Finally, whether the quantitative depletion of GMPs and the qualitative impairment in fate bias arise from a shared upstream mechanism remains an open question.

In conclusion, this study identifies ZNF687 as a regulator of BM myeloid homeostasis that controls both the availability and differentiation competence of osteoclast progenitors. The opposite phenotypes observed in *Zfp687*-KO and *Zfp687*^P937R^-KI models position ZNF687 dosage as a determinant of osteoclastogenic myeloid output in bone.

Together, these findings provide a revised framework for severe ZNF687-related PDB, in which pathological bone remodelling arises not only from altered mature osteoclast function, but also from earlier changes in haematopoietic progenitor dynamics and BM niche support.

## Material and Methods

### Generation of the *Zfp687* knock-out mouse model

To generate the *Zfp687* KO mouse model, we adopted the homologous recombination strategy. A BAC library was used as a template to amplify the *Zfp687* locus for the homologous recombination. PCR fragments were cloned into the pFlrt vector as long, intermediate, and short homology arms, in order to obtain the conditional *Zfp687* allele by homologous recombination using the targeting vector carrying LoxP sites flanking the protein-coding region of exons 2 and 3 of the *Zfp687* gene (**Fig. S1A**). Targeting of the construct was done into the E14Tg2a embryonal stem (ES) cell line, and targeted ES cell clones were identified by Southern blot analysis using a 3’ probe and an internal Neo probe on SphI-cut genomic DNA, and a 5’ probe on BamHI-cut genomic DNA. Sanger sequencing confirmed the presence of the LoxP sites flanking exons 2 and 3 of the *Zfp687* gene. One *Zfp687*-mutated ES clone was injected into C57BL6 blastocysts to establish a mutant *Zfp687*-conditional KO mouse colony. The neomycin resistance gene was removed crossing the mice with a *Flippase*-dependent transgenic mouse. *Zfp687*^FloxP/WT^ mice were then crossed with an *Hprt*-Cre deleter mouse to achieve ubiquitous *Cre*-mediated recombination and germline excision of the floxed allele, generating the constitutive *Zfp687*-KO mouse model (**Fig. S1B**). We performed PCR-based genotyping to confirm the deletion of the *Zfp687* gene using two pairs of allele-specific primers. The genotype distribution followed the expected Mendelian ratios for wild type (WT), heterozygous (*Zfp687*^+/−^), and homozygous (*Zfp687*^−/−^) (**Fig. S1C**), indicating that *Zfp687* depletion is not embryonically lethal. Animals were handled in accordance with the authorization no. 125-2021-PR released by the Italian Ministry of Health; all mice were housed in a pathogen-free barrier environment.

### Body weight and length measurements

Body weight and length were monitored weekly from postnatal day 7 to 3 months of age in WT, heterozygous (*Zfp687*^+/−^), and homozygous (*Zfp687*^−/−^) male mice. Body weight was measured using a precision balance. Body length was measured using a digital caliper, defined as the distance from the tip of the nose to the base of the tail with the animal gently restrained in a standardized position. For skeletal measurements, femurs and tibiae length were collected at sacrifice, cleaned of adherent soft tissues, and bone length was determined using the digital caliper by measuring the distance between the proximal and the distal epiphyses.

### Micro-Computed Tomography (μCT) analysis

Mice were sacrificed by CO_2_ inhalation at the indicated age. The skin was removed, and femurs, tibiae, spine, and skull were cleaned from adherent and soft tissue, fixed in 4% paraformaldehyde, PFA (Sigma-Aldrich, #158127) for 24 h at 4 °C, and then stored in 70% ethanol. μCT analyses were conducted using the SCANCO Medical-μCT40 (Scanco Medical AG, Bassersdorf, Switzerland). For bone morphometry, femoral trabecular and cortical bone were scanned using the following parameters: E=70 kV; I=114 μA; integration time of 600 ms; 6 μm isotropic voxel size. Femurs were scanned 1 mm from the distal board of the growth plate. Cortical thickness was measured at the midshaft region of the femur diaphysis. To compute average cross-sectional area measurement, the volume of interest was divided by the number of slices and voxel height (Tt.Ar=Tt.V/no. of slices x voxel height). Cortical parameters, including total area (Tt.Ar), marrow area (Ma.Ar), and cortical area (Ct.Ar), were calculated according to standardized guidelines for rodent bone microstructure analysis as described by Bouxsein et al^62^. For femur trabecular bone and for midshaft cortical bone, 209 and 36 slices were analysed, respectively. For trabecular bone of femur, the structural parameters bone volume/total volume (BV/TV), trabecular thickness (Tb.Th), trabecular number (Tb.N), and trabecular separation (Tb.Sp) were considered. To reconstruct solid 3D images, selected bone samples were scanned in high resolution. Femur samples were scanned using the following parameters: E=70 kV; I= 14 μA; integration time of 300 ms; 6 μm isotropic voxel size. The reconstructed solid 3D images were applied for visualizing bone morphology and microarchitecture.

### Histological analysis

For histology, femur samples were decalcified in 14% ethylenediaminetetraacetic acid, EDTA (Sigma-Aldrich, #27285) for 14 days, replacing the solution every 3 days. Then, bone samples were dehydrated with ethanol series (70, 85, 95, 100% Et-OH), treated with xylene, and paraffin-embedded. Bone slices of 3 μm were obtained by manual microtome. Bone sections were subjected to wax-removal procedure by xylene treatment and rehydration through a graded series of alcohol (100, 90, 80, 70% Et-OH) and tap water. Bone sections were stained with tartrate-resistant acid phosphatase (TRAP) (Sigma-Aldrich, #387A) to detect osteoclasts activity, according to standard protocols. After staining, bone sections were dehydrated, mounted with the mounting medium (Bio-Optica, #05-BMHM), covered with a coverslip, and analysed by transmission light microscopy, using Nikon Motorized Eclipse Ni-U Microscope and Nikon Manual Optical Microscope. Osteoclasts surface and number were measured by the TRAPHisto open-source software^63^.

### Cell culture and differentiation

Primary murine bone marrow monocytes/macrophages (BMMs) were obtained from the bone marrow of 12 week-old, male mice. Briefly, euthanized through CO_2_ and immediately after the sacrifice, femurs and tibiae were carefully cleaned of all the connective tissues; both the distal and proximal ends of bones were cut, and the BM was centrifuged out. After centrifuge, total BM cells were cultured in MEM α, nucleosides medium (Gibco, 22571120) comprising 10% FBS, 1% penicillin/streptomycin, and 1% L-Glutamine (complete MEM α), supplemented with 100 ng/mL hM-CSF (PeproTech - 300-25) at 37°C, 7% CO_2_. After 5 days of culture, the cells were washed gently with PBS, and the adherent cells were harvested as BMMs for further experiments. For osteoclast differentiation, cells were detached using TrypLE Express Enzyme (1X) (Gibco, 12605010), and seeded in 96-well at a density of 15×10^3^ cells per well in triplicate in complete MEM α, supplemented with 30 ng/ml hM-CSF and 150 ng/ml sRANKL (PeproTech, 315-11). Cell media was replaced every 2 days. After 5-7 days of culture, cells were washed 2X with PBS, and fixed with 4% paraformaldehyde (PFA) for 20 min, then stained with tartrate-resistant acid phosphatase (TRAP) (Sigma-Aldrich, 387A). Osteoclast area and osteoclast-covered surface area quantifications were determined using ImageJ software.

RAW264.7 cells were cultured in DMEM High Glucose GlutaMAX (Gibco, 10569010), with 10% FBS, 1% penicillin/streptomycin, and 1% L-Glutamine at 37 °C, 5% CO_2_. For osteoclast differentiation, 5×10^3^ cells were plated in 24-well plates and the medium was switched in MEM α GlutaMAX (Gibco, 32561037), with 10% FBS, 1% penicillin/streptomycin, and 1% L-Glutamine. The day after, medium was changed and supplemented with 100 ng/ml sRANKL for the osteoclastogenic induction without or with 30 ng/ml hM-CSF. The medium was changed every 48 h, until the end of the differentiation (5 days upon stimulation). Differentiated osteoclasts were fixed in 4% PFA and stained with tartrate TRAP (Sigma-Aldrich #387A).

For proliferation assay, RAW264.7 cells were seeded at a density of 3×10^3^ cells per well in triplicate in MEM α GlutaMAX (Gibco, 32561037), with 10% FBS, 1% penicillin/streptomycin, and 1% L-Glutamine, without or with 30 ng/ml hM-CSF. The day after, medium was changed and supplemented with Cell Counting Kit-8 reagent (Dojindo, #CK04), according to manufacturer’s instructions. Absorbance was measured at 450 nm using the microplate reader PerkinElmer luminometer (Victor X3) at the indicated time points (3, 6, 12, 24, 48, and 72 hours). Proliferation values were normalized to the 3-hour time point, which was used as the baseline reference.

Primary murine BM-derived mesenchymal stromal cells (BM-MSCs) were obtained from femurs and tibiae of 8-weeks-old mice. Briefly, mice were euthanized through CO_2_ and immediately after the sacrifice, femurs and tibiae were carefully cleaned of all the connective tissues; both the distal and proximal ends of bones were cut, and the BM was centrifuged out. ACK (Ammonium-Chloride-Potassium) lysing buffer (Gibco, A1049201) was used to eliminate red blood cells. Total BM cells were cultured in complete expansion medium (MesenCult Expansion Kit Mouse #05513, Stem Cell Technologies), at 37 °C, 7% CO_2_. BM-MSCs were expanded for 7 days. For the osteogenic differentiation, cells were detached using 0.25% Trypsin-EDTA, plated in 24-well plates and cultured in complete expansion medium until they reached 80-90% confluency. Then, to induce osteogenic differentiation, medium was replaced with complete MesenCult Osteogenic Medium (MesenCult Osteogenic Stimulatory Kit Mouse #05504, Stem Cell Technologies), and cells were cultured at 37 °C, 7% CO_2_. Medium was changed every 3 days for 8 days.

### Protein extraction and Western blotting

Total protein extraction from murine cell lines (RAW264.7 cells and BMMs) was performed in RIPA buffer (Thermo Scientific, #89900) with 1X proteinase inhibitor cocktail (Applied Biological Materials #G135) and 1X phosphatase inhibitors cocktail (Sigma-Aldrich, #P0044). Protein quantification was obtained by the BCA protein assay kit (Thermo Scientific, #23227). Protein samples were boiled at 85 °C for 3’ and then were separated by SDS-PAGE electrophoresis, using 8-16% Tris-Glycine gels (Invitrogen #XP08160). Samples were transferred on a nitrocellulose membrane (Invitrogen #IB23002), blocked with 4% w/v non-fat dry milk dissolved in TBS-T (1X TBS, 0.01% Tween-20) for 1 h at RT. Primary antibodies used for the Western blot experiments were rabbit anti-BRD4 (1:1000, CST #83375), rabbit anti-c-Myc (1:1000, CST #5605), rabbit anti-NFAT2 (1:1000, CST #8032), and rabbit anti-Vinculin (1:1000, CST #13901). Membranes were incubated with secondary antibodies conjugated with HRP for 1 h at RT. The bands were visualised using enhanced chemiluminescence detection reagents (SuperSignal West Pico PLUS Chemiluminescent Substrate, Thermo Scientific #34577). Equal loading was confirmed by using antibody against anti-Vinculin. The intensity of the western blot signals was determined by densitometry analysis using the ImageJ software and normalised to the density value of the loading control.

### RNA isolation, qRT-PCR, and ddPCR analysis

Total RNA extraction (from BM-MSCs, mouse osteoblasts, BMMs, and RAW264.7 cells) was obtained using TRI-Reagent (Sigma-Aldrich #T9424), following the manufacturer’s instruction. One microgram of total RNA was retrotranscribed in cDNA using the LunaScript RT SuperMix kit (NEB, E3010). qRT-PCR was performed using the SYBR Select Master Mix for CFX (Applied Biosystems) and specific primers on CFX Opus RT PCR System instrument. The transcript levels were normalized to the levels of *Hprt* within each sample, and the ΛλΛλCT method was used. The reaction was conducted in triplicate.

For the ddPCR, the Bio-Rad QX200 Droplet Digital PCR Systems (Bio-Rad) was used following the manufacturer instructions. In detail, a volume corresponding to 5 ng of cDNA template was added to the QX200 ddPCR EvaGreen Supermixmaster (Bio-Rad, #1864034). Then, the reaction mix underwent droplet generation by the QX200 droplet generator (Bio-Rad). Droplet-partitioned samples were then transferred to a 96-well plate, sealed and amplified to thermal cycler. Plates containing amplified droplets were loaded into the QX100 droplet reader (Bio-Rad). QuantaSoft Software (Bio-Rad) and QX Manager Software (QX Manager Standard Edition Version 2.1, Bio-Rad) were used to collect and analyse the dataset obtained by ddPCR reaction. A single manual fluorescence threshold was applied across all wells, based on negative controls and cluster separation.

### Cell Isolation for FACS and flow cytometry analyses

Long bones (femurs and tibias) were collected from 3-month-old male mice as described above. The BM was centrifuged out, and ACK lysing buffer was used to remove red blood cells. Prepared single-cell suspensions (2×10^6^ BM cells) were washed with ice-cold FACS buffer (0.5% BSA + 1 mM EDTA in 1X PBS) and incubated with Fc blocking buffer (Miltenyi Biotech, #130-092-575) for 20 min at 4 °C. Cells were incubated in the dark for 30 min at 4 °C with primary antibody solution and washed 2 times with FACS buffer, and re-suspended in FACS buffer with 7-AAD Viability Staining Solution (Sony Biotechnology, #2702020). FACS was performed using a Becton Dickinson Aria II equipped with 3 lasers (BD Bioscience). Fluorescence minus one (FMO) and Isotypes (Iso) controls were used for additional compensation and to assess background levels for each stain. Gates were drawn as determined by internal FMO controls to separate positive and negative populations for each cell surface marker. Typically, 1 million events were recorded for each FACS analysis, and the data were analysed using FlowJo (v10.10.0).

### FACS antibodies

Antibodies for FACS of BM samples included Ter119 (clone Ter-119, Sony Biotechnology #1181100), B220 (clone RA3-6B2, Sony Biotechnology #1116145), CD117 (clone 2B8, Sony Biotechnology #1129040), CD115 (clone AFS98, Sony Biotechnology #1277620), CD11b (clone M1/70, Sony Biotechnology #1106025), CD45 (clone 30-F11, Sony Biotechnology #1115650), F4/80 (BM8, Sony Biotechnology #1215580).

### scRNA-sequencing

For scRNA-seq libraries we sorted 7-AAD^−^Ter119^−^B220^−^CD117^+^ cells from BM of n=2 WT and n=2 *Zfp687*^−/−^ mice at 12 weeks of age. BM cells were isolated and prepared for FACS sorting as described above. scRNA-sequencing libraries were prepared using the 10x Genomics Chromium Single Cell 3’ v4 (polyA) chemistry, targeting approximately 10000 cells per sample.

### scRNA-seq data pre-processing

Sequencing was performed on Illumina. Raw sequencing reads were aligned to the mouse reference genome GRCm39 (annotation version GRCm39-2024-A) using CellRanger v9.0.1 with intronic reads included.

Downstream analysis was performed in R using Seurat v5. For each sample independently, cells were filtered based on the following quality control criteria: number of detected genes between 500 and 6’000, total UMI count greater than 1000, and mitochondrial gene fraction below 10%. Doublets were identified and removed using scDblFinder. Mitochondrial genes and *Malat1* were excluded from the feature set prior to normalization; sex-linked genes were retained given the same-sex, same-litter experimental design. After filtering and doublet removal, 30600 high-quality singlets were retained across the four samples.

### Integrated analysis of single cell datasets

Normalization was performed independently per sample using SCTransform v2 (vst.flavor = “v2”), which fits a regularized negative binomial regression model to account for sequencing depth differences. Cell cycle scoring was performed using the cc.genes.updated.2019 gene sets with gene names converted to title case for compatibility with the mouse genome. To preserve biologically meaningful cycling versus quiescent cell distinctions, particularly relevant in the BM hematopoietic context, partial cell cycle regression was applied by regressing the difference between S phase and G2/M phase scores (CC.Difference = S.Score - G2M.Score) during SCTransform, following the approach described by Tirosh et al^64^. This retains the distinction between actively cycling and quiescent progenitors while removing intra-cycle phase variability. Principal component analysis (PCA) was performed on the SCT-normalized data using the top 3000 variable features, retaining 50 principal components. Technical variation across samples was corrected using Harmony (theta=2, correcting for sample identity), and the first 20 Harmony-corrected dimensions, selected based on elbow plot inspection, were used for UMAP embedding and graph-based clustering. Shared nearest neighbor (SNN) graph construction was performed with FindNeighbors (dims=1:20), and clustering was performed with FindClusters at resolution 0.5, yielding 22 transcriptionally distinct clusters.

### Visualization and clustering

Cluster annotation was performed using a two-step approach. First, canonical lineage marker expression was visualized by DotPlot for each cluster using a curated panel of myeloid, monocytic, erythroid, and progenitor markers. Second, label transfer from the Paul et al. 2015 Lin⁻Kit⁺ murine bone marrow dataset (GSE72857; 2730 cells, 19 annotated clusters^41^) was performed using FindTransferAnchors and TransferData with SCTransform normalization, applying the label transfer to all cells simultaneously to avoid genotype-specific bias. Final cluster annotations were assigned manually, prioritizing canonical marker expression over label transfer predictions, particularly for clusters with prediction scores below 0.5 or for primitive progenitor populations absent from the Paul 2015 reference^41^.

### Reconstructing cell development trajectories

Trajectory inference and pseudotime analysis were performed using Monocle3. The Seurat UMAP embedding and Harmony-corrected PCA dimensions were transferred to the Monocle3 cell_data_set object. The principal graph was learned on the full dataset without partition constraints (use_partition = FALSE), and pseudotime was calculated by rooting the trajectory at the HSC/MPP cluster. Cell cycle phase composition within individual clusters was compared between genotypes using Fisher’s exact test on the proportion of G1 versus cycling (S + G2M) cells.

All analyses were performed in R v4.6.0. Key package versions: Seurat v5, harmony v2.0.2, scDblFinder, speckle, clusterProfiler, monocle3.

### CellRank fate probabilities

Lineage fate probabilities were computes using CellRank (v2.2.0)^44^. A combined kernel was constructed as a weighted sum of PseudotimeKernel (weigth 0.8), based on Monocle3 pseudotime, and a ConnectivityKernel (weight 0.2), capturing transcriptional similarity. Terminal states were identified using the GPCCA estimator (Generalized Perron Cluster Cluster Analysis), and fate probabilities toward each microstate were computed. Monocyte/macrophage fate probability was defined as the sum of the probabilities toward all monocyte- and macrophage-associated terminal states, excluding dendritic cell (DC) states (“narrow” definition). Fate probabilities were compared between genotypes both across all cells and restricted to the Early GMP cluster, using Wilcoxon rank-sum test, with effect sizes reported as rank-biserial correlation (r).

### Generation of hiPSCs and hematopoietic differentiation

Human dermal fibroblasts (hDFs) were obtained by skin biopsy from a Paget’s disease of bone patient carrying the ZNF687 P937R mutation and from his unaffected sibling, who served as a healthy control with similar genetic background. hDFs were cultured in DMEM high Glucose GlutaMAX (Gibco, 10569010), supplemented with 20% FBS. Cells were maintained at 37°C, 5% CO_2_. For episomal reprogramming nucleofection was performed using the Human Dermal Fibroblast Nucleofector Kit (NHDF-Adult, Cat. No. VPD-1001), according to manufacturer’ instructions. Briefly, approximately 1×10^6^ cells were resuspended in the nucleofection solution and mixed with the episomal plasmid combination: pCXLE-hOCT3/4-shp53-F (Addgene, #27077), pCXLE-hSK (Addgene, #27078), and pCXLE-hUL (Addgene, #27080). Immediately following nucleofection, cells were resuspended in warm culture medium and plated. Twenty-four hours post-nucleofection (Day1), the culture medium was refreshed. On Day 2, transfected cells were re-seeded onto a Matrigel-coated 6-well plate. Starting from Day 3, the culture medium was transitioned to DMEM F12 with Hepes supplemented with the N2B27 medium, made of N2 Supplement, B27 Supplement, 100 µg/ml of bGFG, β-mercaptoethanol, and 10 mM of MEM non-essential amino acid solution. On Day 8, the N2B27 medium was replaced by mTeSR1. On Day22, individual hiPSCs colonies were mechanically picked and transferred onto fresh matrix-coated plates for further expansion and characterization.

### Immunostaining of iPSCs

Selected iPSCs of healthy and P937R-mutant donors were validated for pluripotency by immunofluorescence staining for the pluripotency markers NANOG and OCT3/4. Briefly, iPSCs were fixed in 4% PFA for 15’ and subsequently and permeabilized using a blocking solution containing 10% FBS, 1% BSA, and 0.5% Triton X-100 in 1X PBS for 30’ at RT. Primary antibodies used for the immunostaining were mouse OCT-3/4 (C-10) (sc-5279; 1:400), rabbit NANOG (D73G4) (CST 4903; 1:500). Following secondary antibodies incubation (Goat anti-Mouse AF546, Invitrogen A11030, and goat anti-rabbit AF488, Invitrogen A11008; 1:400), cells were counterstained with DAPI. Images were acquired by Leica DMI 6000 microscope.

### Haematopoietic differentiation of iPSCs

ZNF687 P937R and control iPSCs were differentiated into hematopoietic progenitor cells (HPCs) using the STEMdiff Hematopoietic Kit (StemCell Technologies, Catalog #05310). Briefly, one day before initiating differentiation, hiPSCs at 70-80% confluency were treated with Cell Dissociation Buffer to generate aggregates of 100-200 μm in diameter. A total of 100-150 clusters were transferred into Geltrex-coated 6-well plates and cultured overnight. On day 1, StemMACS iPS-Brew-XF medium was replaced with Basal Medium supplemented with Supplement A to initiate mesoderm induction. On day 2, a half-medium change was performed. On day 3, Basal Medium with Supplement A was replaced with Basal Medium supplemented with Supplement B to promote hematopoietic specification. Half-medium changes were subsequently performed on days 5, 7 and 10. On day 12, floating cells were harvested and processed for downstream analyses.

### FACS analysis of HPCs

Immunophenotypic characterization of HPCs was performed by flow cytometry at the end of the differentiation protocol (day 12). Harvested cells were stained with the following primary monoclonal antibodies: anti-CD43 PE (clone eBio84-3C1, Invitrogen #12-0439-42), anti-CD45 FITC (clone HI30, Sony Biotechnology #2120020) and anti-CD34 APC (clone 581, Sony Biotechnology #2317550). Isotype controls were performed using monoclonal mouse IgG1 kappa isotype control antibodies conjugated to PE (clone MOPC-21, Sony Biotechnology #2600700), FITC (clone MOPC-21, Sony Biotechnology #2600690), and APC (clone MOPC-21, Sony Biotechnology #2600710). Live cells were gated based on FSC/SSC parameters. Acquisition was performed on a FACSAria III system (BD Biosciences), collecting a minimum of 10000 events per sample. Data were analysed using FlowJo (v10.10.0).

### CFU-C assay

To assess the hematopoietic colony-forming potential of the generated HPCs, cells were plated in MethoCult H4434 Classic methylcellulose-based medium (#04434, StemCell Technologies) according to the manufacturer’s instructions. Colony formation was monitored at days 12 and 15, with day 15 representing the end of the differentiation. Colonies were counted and classified by morphology into the following categories: colony-forming unit granulocyte-macrophage (CFU-GM), burst-forming unit erythroid (BFU-E), colony-forming unit erythroid, and colony-forming unit granulocyte-erythroid-macrophage (CFU-GEMM). At the end of the assay, representative images of each colony type were acquired at 5× magnification using an inverted DMI6000 microscope (Leica Microsystems).

### Quantification and statistical analyses

All data are presented as mean or median ± standard deviation (s.d.). The sample size for each experiment and the replicate number of experiments are included in the figure legends. Statistical significance was defined as p<0.05. Statistical analyses were performed using ordinary one-way ANOVA, student’s-T-test, and multiple unpaired t test(GraphPad Prism; version 9.3.1). All single-cell RNA-sequencing data analyses and relative statistical analysis were performed in R v4.6.0.

## Acknowledgments.

Authors acknowledge members of the “Bone Diseases and Tumors” laboratory at IGB-CNR for constructive feedback provided during manuscript preparation. We are grateful to members of the Integrated Microscopy and FACS Facilities of IGB-CNR.

The research leading to these results has received funding from the Italian Association for Cancer Research (AIRC; project ID 25110) to F.G., by the Ministry of University and Research under the PRIN 2022 PNRR Call (project code P20224JCNN) to F.G.; by the Next Generation EU programme within the framework of the National Recovery and Resilience Plan (PNRR), Investment PE8 – Project Age-It, to F.G.; and by the MARADONA Project, funded under the Campania Region Cohesion Agreement “Rare Diseases” Call.

## Author contributions

S.R. and F.G. conceived and designed the study. S.R. performed the majority of the experiments. F.G. supervised the study and data analysis. A.M. contributed to the osteoclastogenesis study with RAW264.7 cell line. D.L. performed the initial analysis of the scRNA-seq data. C.S. contributed to the μCT study and data analysis. D.A. and A.S. generated the *Zfp687*-KO mouse model. V.L., M.S. and M.M. performed iPSC differentiation into haematopoietic and myeloid precursors. All authors read and approved the final version of the manuscript.

## Data availability

All relevant data supporting the key findings of this study are available within the article.

---

**Supplementary Figure 1:**
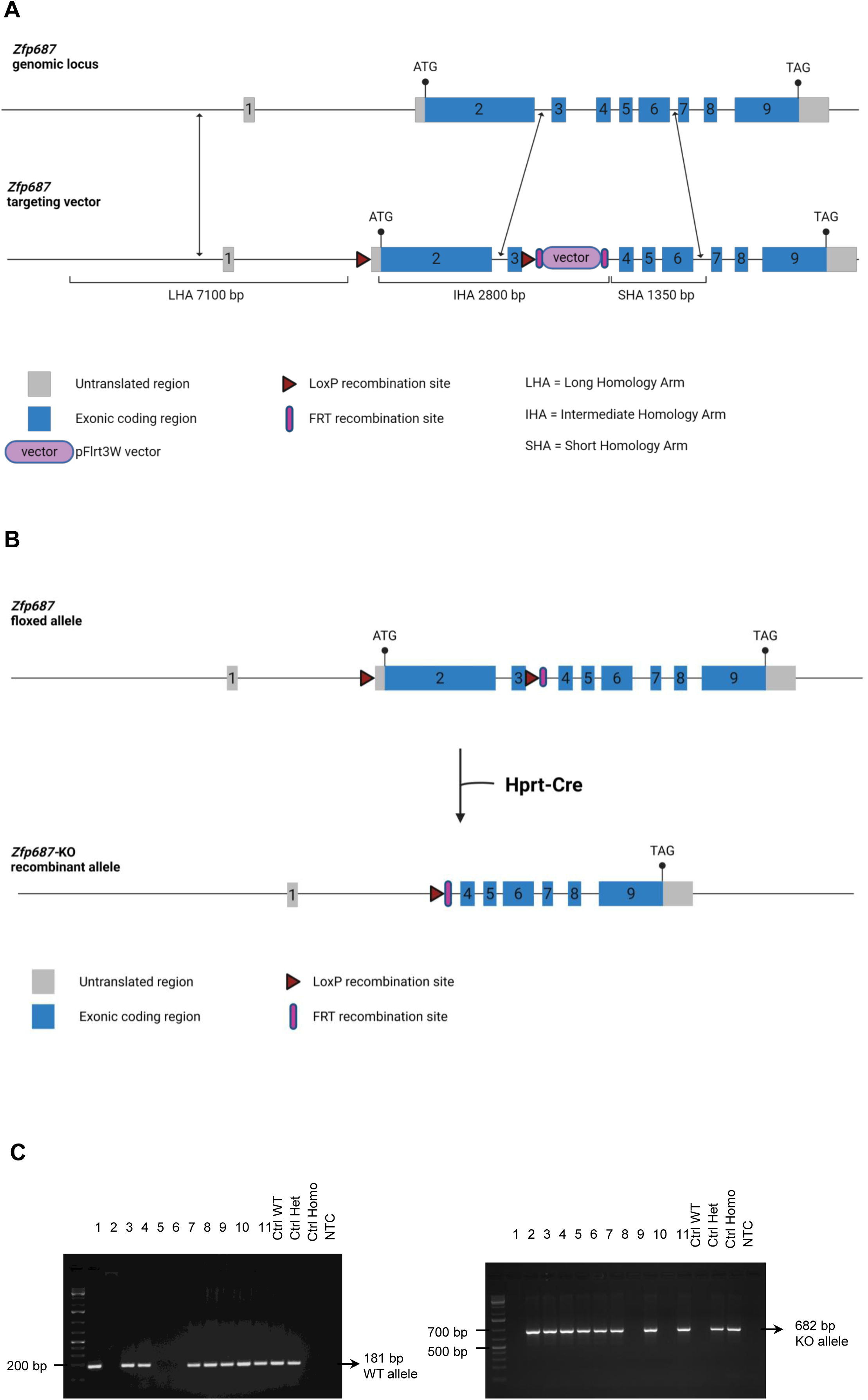
**A)** Schematic overview of the *Zfp687* targeted allele (up) and the construction of the targeting vector (down) for the *Zfp687*-KO mouse model generation. Combination of FRT and LoxP sites allows manipulation of targeted allele in mice. **B)** Conditional mice were crossed with germline *Hprt*-Cre mice to generate constitutive *Zfp687*-KO mice. Exons are indicated with boxes; grey boxes indicate untranslated regions; light-blue boxes indicate coding regions. The long homology arm of the targeting vector includes a genomic region of 7077 bp, the intermediate homology arm one of 2799 bp, and the short homology arm one of 1379 bp. The exons 2 and 3 are flanked by LoxP sites, and the Neo cassette (pFlrt3W vector) is flanked by FRT sites. **C)** Representative agarose gels showing PCR-based genotyping of *Zfp687*-KO mice. (Left) PCR amplification of the WT *Zfp687* allele across 11 animals (lanes 1-11). Lanes 12-14 show positive control reactions for WT, heterozygous (*Zfp687*^+/−^), and homozygous KO (*Zfp687*^−/−^) genotypes, respectively. Lane 15 shows a no-template negative control (NTC). The WT allele yields an amplicon of 181 bp. (Left) PCR amplification of the KO *Zfp687* allele across the same 11 animals (lanes 1-11), with identical positive controls for WT, *Zfp687*^+/−^, and *Zfp687*^−/−^ genotypes (lanes 12-14) and a NTC (lane 15). The KO allele yields an amplicon of 682 bp.

**Supplementary Figure 2:**
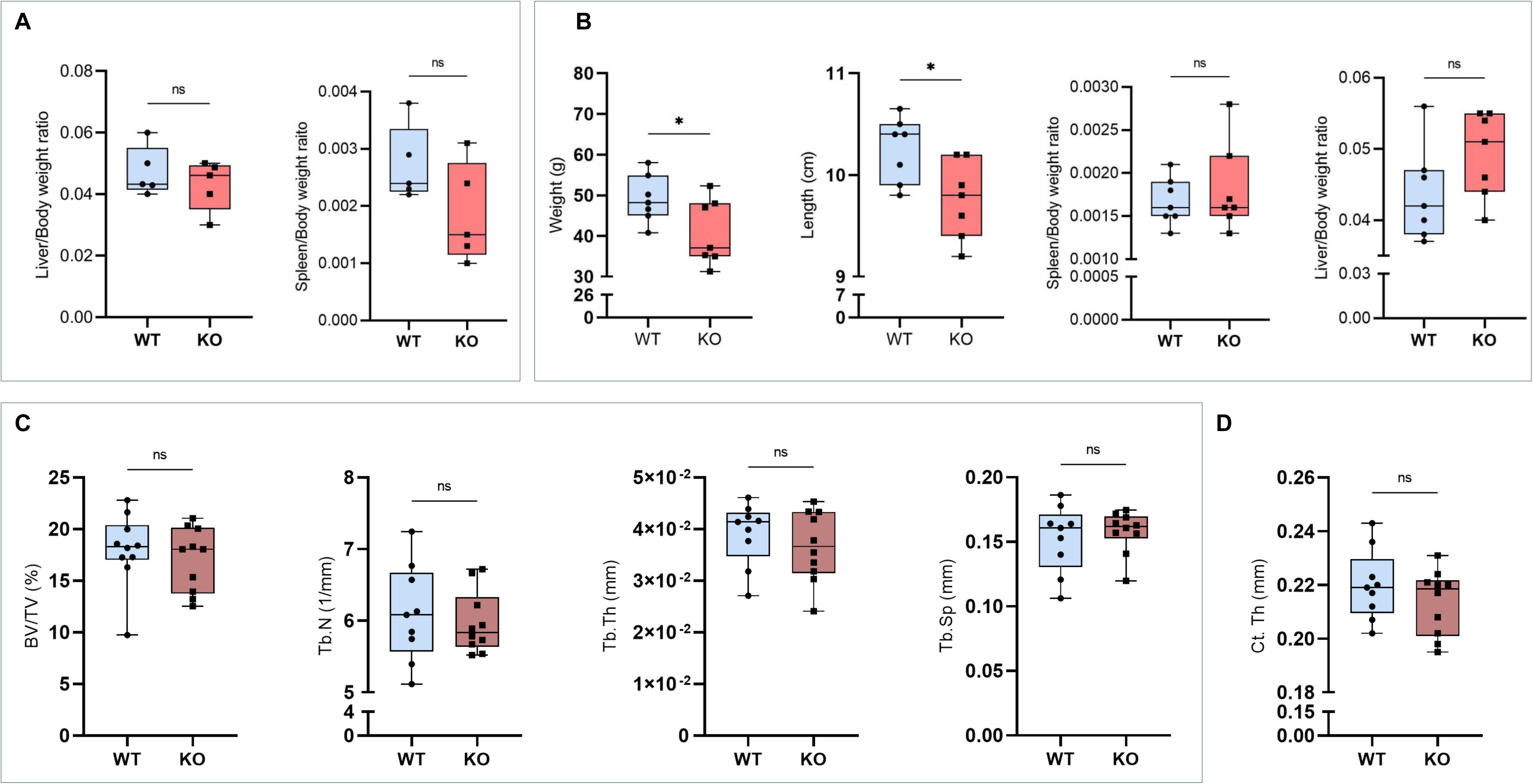
**A)** Bar graph showing (left) liver- and (right) spleen-to-body weight ratio in 3-month-old WT (n=5) and KO (n=5) mice. Data are presented as median±s.d. Statistical significance was assessed by two-tailed unpaired t test (ns=not significative). **B)** Measurements of body weight, length, liver- and spleen-to-body weight ratio of 8-month-old WT and KO (n=7) mice. Data are shown as mean±s.d. Statistical significance was assessed by two-tailed unpaired t test (ns=not significative; *p<0.05). **C)** Trabecular bone parameters, including BV/TV, Tb. N, Tb. Th, and Tb. Sp determined by µCT in femoral distal epiphysis of 3-month-old WT and KO (n=10) mice. Data are presented as median±s.d. and statistical significance was assessed by two-tailed unpaired t test (ns=not significative). **E)** Cortical bone parameter (Ct. Th) was determined by µCT scan in the femoral midshaft of 3-month-old WT and KO (n= 9) mice. Data are presented as median±s.d. and statistical significance was assessed by two-tailed unpaired t test (ns=not significative).

**Supplementary Figure 3:**
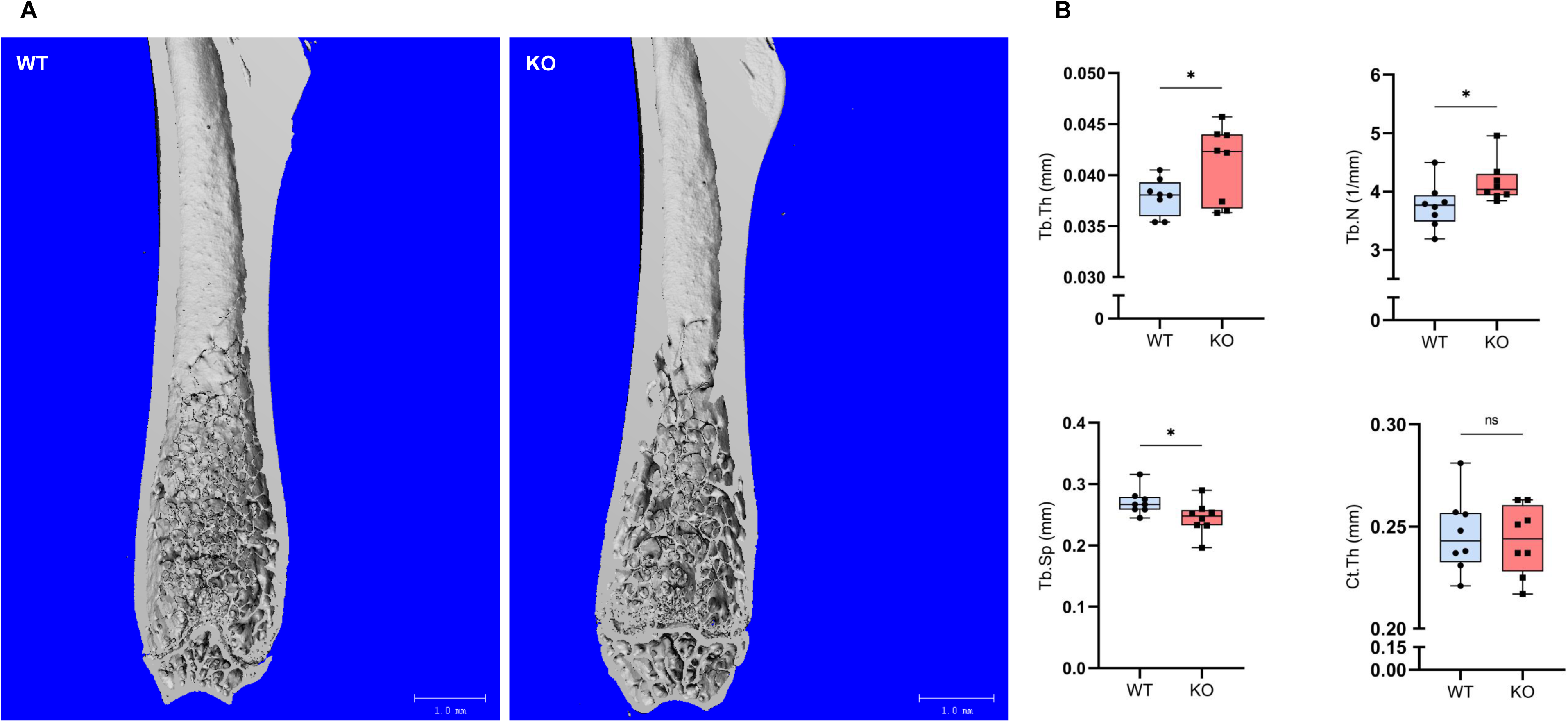
**A)** Representative µCT 3D reconstruction of femurs from 8-month-old WT and KO mice. Scale bars: 1 mm. **B)** Trabecular bone parameters, including Tb.Th, Tb.N, and Tb.Sp, and Ct.Th were determined by µCT scan in femoral distal epiphysis and in the femoral midshaft of 8-month-old WT and KO (n=8) mice. Data are presented as median±s.d. and statistical significance was assessed by two-tailed unpaired t test (ns=not significative; *p<0.05).

**Supplementary Figure 4:**
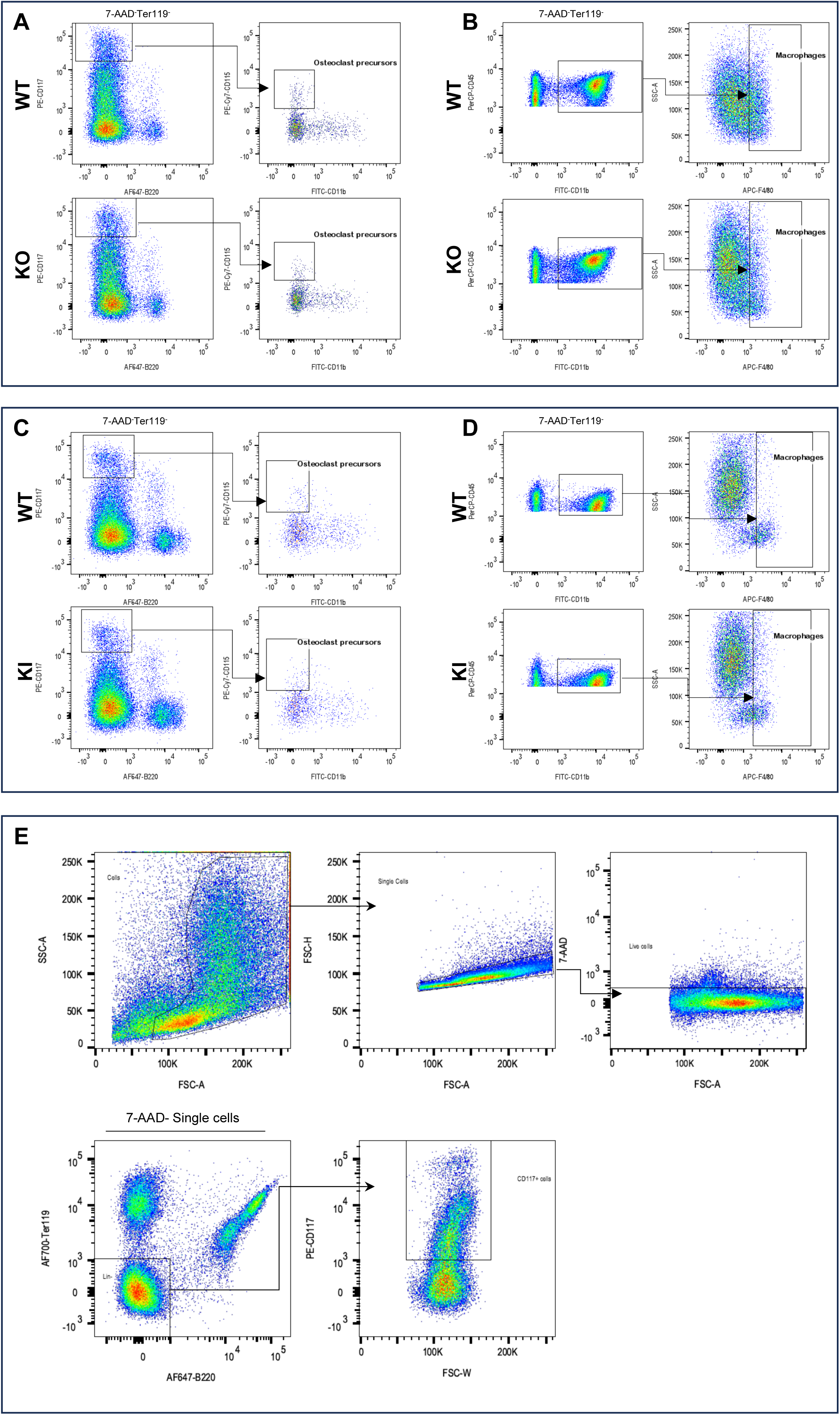
**A)** Representative pseudocolor plots showing the gating strategy on 7-AAD^−^Ter119^−^ cells to evaluate BM OC precursors frequency (CD117^+^B220^−^CD115^+^CD11b^−^) in 3-month-old WT and KO mice. **B)** Representative pseudocolor plots showing the gating strategy on 7-AAD^−^Ter119^−^ cells to evaluate BM Mφ (CD45^+^CD11b^int/high^F4/80^+^) frequency in WT and KO. **C-D)** Representative pseudocolor plots showing the gating strategy as described in **A-B)** for WT and KI mice. **E)** Representative FACS gating strategy to sort 7-AAD^−^Ter119^−^B220^−^CD117^+^ cells from WT and KO BM-derived cells.

**Supplementary Figure 5:**
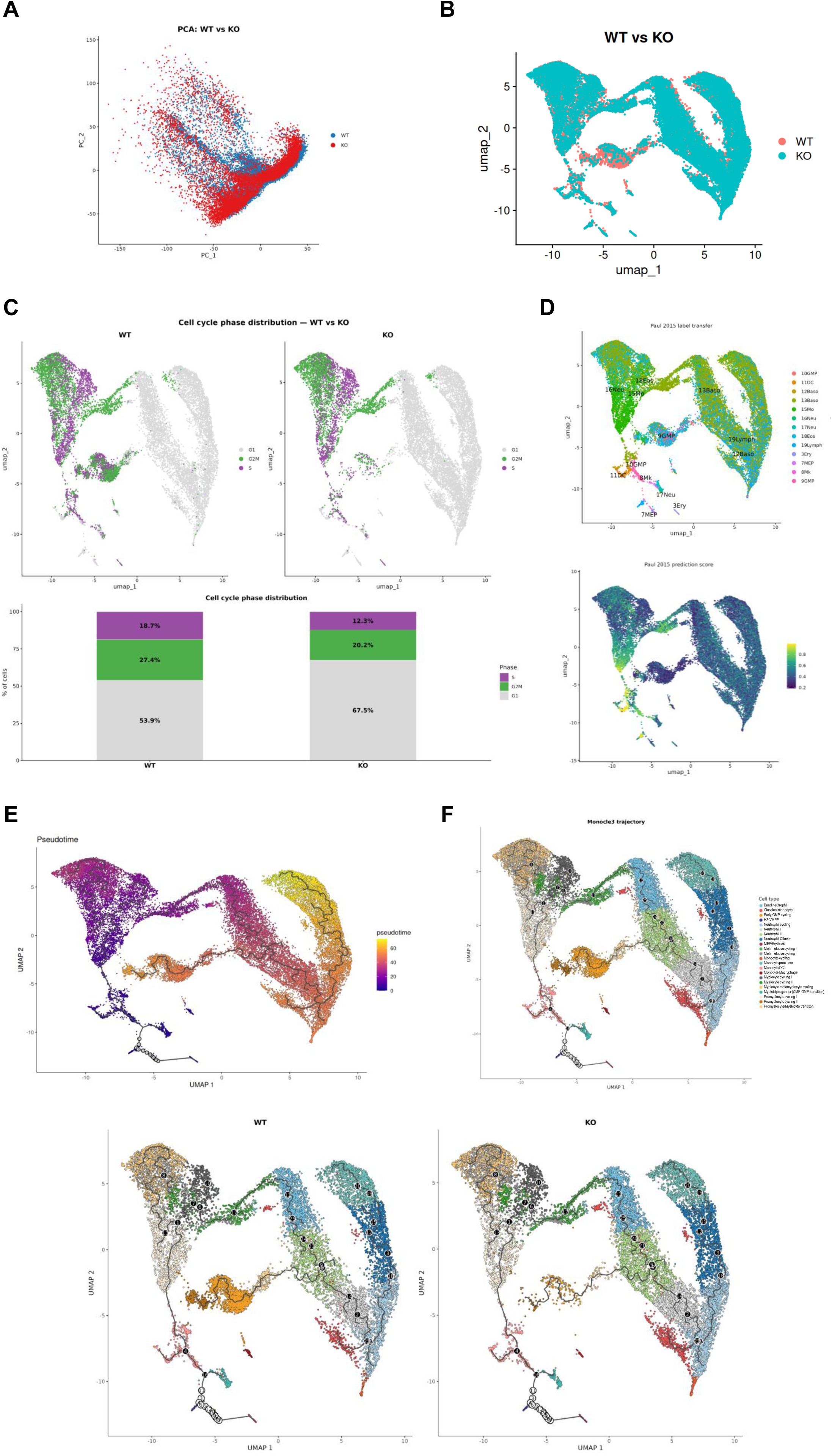
**A)** PCA of the 4 samples prior to Harmony batch correction. Cells are coloured by genotype (WT, blue; KO, red), illustrating the pre-integration genotype distribution in PCA space. **B)** UMAP visualization of all 30600 singlets after Harmony batch integration, coloured by genotype (WT, pink; KO, light-blue). **C)** Top: UMAP split by genotype showing cell cycle phase assignment (G1, grey; S, purple; G2M, green) prior to partial cell cycle regression. Bottom: stacked bar plot quantifying the percentage of cells in each cell cycle phase per genotype. Proportions are comparable between WT (G1 53.9%, S 18.7%, G2M 27.4%) and KO (G1 67.9%, S 17.2%, G2M 14.9%. **D)** Top: UMAP coloured by Paul 2015 predicted cell type labels^41^, shown in the original nomenclature. Bottom: UMAP coloured by Paul 2015 prediction score (range 0-1), reflecting the confidence of label transfer for each cell. **E)** Ridgeline plot showing UMAP pseudotime density distribution per annotated cluster. The learned trajectory (black line) recapitulates the myeloid differentiation continuum, with pseudotime values, ranging from 0 (HSC/MPP root) to >60 (yellow, terminally differentiated cells). **F)** Monocle3 principal graph trajectory overlaid on the UMAP embedding all cells, coloured by annotated cell type. Numbered black circles indicate branch points. The trajectory is rooted at HSC/MPP cluster. Split trajectory visualization between WT and KO mice is displayed at the bottom.

